# Mouse Brain Extractor: Brain segmentation of mouse MRI using global positional encoding and SwinUNETR

**DOI:** 10.1101/2024.09.03.611106

**Authors:** Yeun Kim, Haley Hrncir, Cassandra E. Meyer, Manal Tabbaa, Rex A. Moats, Pat Levitt, Neil G. Harris, Allan MacKenzie-Graham, David W. Shattuck

## Abstract

In spite of the great progress that has been made towards automating brain extraction in human magnetic resonance imaging (MRI), challenges remain in the automation of this task for mouse models of brain disorders. Researchers often resort to editing brain segmentation results manually when automated methods fail to produce accurate delineations. However, manual corrections can be labor-intensive and introduce interrater variability. This motivated our development of a new deep-learning-based method for brain segmentation of mouse MRI, which we call Mouse Brain Extractor. We adapted the existing SwinUNETR architecture (Hatamizadeh et al., 2021) with the goal of making it more robust to scale variance. Our approach is to supply the network model with supplementary spatial information in the form of absolute positional encoding. We use a new scheme for positional encoding, which we call Global Positional Encoding (GPE). GPE is based on a shared coordinate frame that is relative to the entire input image. This differs from the positional encoding used in SwinUNETR, which solely employs relative pairwise image patch positions. GPE also differs from the conventional absolute positional encoding approach, which encodes position relative to a subimage rather than the entire image. We trained and tested our method on a heterogeneous dataset of N=223 mouse MRI, for which we generated a corresponding set of manually-edited brain masks. These data were acquired previously in other studies using several different scanners and imaging protocols and included *in vivo* and *ex vivo* images of mice with heterogeneous brain structure due to different genotypes, strains, diseases, ages, and sexes. We evaluated our method’s results against those of seven existing rodent brain extraction methods and two state-of-the art deep-learning approaches, nnU-Net (Isensee et al., 2018) and SwinUNETR. Overall, our proposed method achieved average Dice scores on the order of 0.98 and average HD95 measures on the order of 100 µm when compared to the manually-labeled brain masks. In statistical analyses, our method significantly outperformed the conventional approaches and performed as well as or significantly better than the nnU-Net and SwinUNETR methods. These results suggest that Global Positional Encoding provides additional contextual information that enables our Mouse Brain Extractor to perform competitively on datasets containing multiple resolutions.

## 1 Introduction

Preclinical models play a key role in the study of biological processes, with the goal of translating knowledge gained from these studies into a better understanding of aspects of human systems. The mouse has been a critical component in many translational studies and has been particularly important in the investigation of the processeses of neurodevelopment and neurodegeneration. Magnetic resonance imaging (MRI) of mouse models provides a mechanism for quantifying effects in preclinical studies, thereby enabling statistical analyses of differences between groups or changes over time. Wherever possible, it is desirable to automate the computational processing of preclinical MRI to reduce operator time required for analysis, to reduce rater-dependent variability, and to facilitate reproducibility.

One preliminary step that is essential in many analysis workflows for mouse brain MRI is the segmentation of brain from non-brain tissue, a process widely-known as brain extraction or skull stripping. The extracted brain-only images can be used directly to study whole brain characteristics and, importantly, can facilitate downstream processing methods. One major application is the use of skull-stripping prior to brain registration, where the isolation of the brain region can improve alignment to another brain image by reducing the spatial domain of interest and removing highly variable non-brain anatomical features that are not relevant to the alignment task.

Although numerous automated methods have been developed for brain extraction in human MRI, fewer tools have been created specifically for rodent imaging. In some cases, researchers have adapted automated tools that were originally developed for processing human MRI, which has led to variants that include SPM-Mouse (Sawiak et al., 2013), based on SPM (Ashburner, 2012), and the Rodent Brain Extraction Tool (rBET; Wood et al., 2013), based on FSL’s Brain Extraction Tool (BET; Smith, 2002). Preclinical researchers also make use of human-specific tools by modifying image metadata to meet a method’s expectations, e.g., by changing the resolution specified in the image file such that the software will interpret its voxels to be larger and thus scale the mouse brain to human proportions. This technique has been used, for example, with ANTs’ antsBrainExtraction (Avants et al., 2011). However, methods originally designed for human MRI may yield poor-quality results in mouse data due to differences in head shape, anatomical structure, and tissue composition.

A small number of tools have been developed specifically for performing brain extraction in rodent MRI. Among these is the graph-based 3D Pulse-Coupled Neural Network (3D-PCNN), which iteratively modifies brain mask regions based on the similarity of image intensities in adjacent voxels (Chou et al., 2011). However, in images with lower tissue contrast or substantial field inhomogeneity artifacts, 3D-PCNN can perform poorly (Oguz et al., 2014). Other approaches include Rapid Automatic Tissue Segmentation (RATS; Oguz et al., 2014) and SHape descriptor selected Extremal Regions after Morphologically filtering (SHERM; Y. Liu et al., 2020). RATS and SHERM both use iterative mathematical morphology operations to form initial regions of interest. RATS follows this step by converting its initial binary mask into a surface and then performing Layered Optimal Graph Image Segmentation for Multiple Objects and Surfaces (LOGISMOS) to improve the topology of the surface (Oguz et al., 2014). In contrast, SHERM analyzes the initial mask to detect maximally-stable extremal regions (Matas et al., 2004) which are clusters of voxels composed of similar grey-scale values surrounded by sharp intensity gradients at the edges. The detected regions are then filtered based on convexity and shape descriptors, and are finally merged to produce a final brain segmentation (Y. Liu et al., 2020). RATS and SHERM are sensitive to image contrasts because they both rely on image intensity gradients to define the edges of the brain. Another disadvantage of 3D-PCNN, RATS, and SHERM is that they often require a manual search for unique parameters to produce optimal results for each image (Y. Liu et al., 2020).

In cases where automated tools fail to produce accurate segmentation results, researchers frequently resort to manual editing of the generated outputs. Manual segmentation is often regarded as a gold standard, but the manual correction process can be labor-intensive and is susceptible to intra-rater and inter-rater variability. In an effort to directly emulate well-edited segmentations, many brain extraction methods have been designed using machine learning methods that are trained on collections of manually-delineated data.

The advent of deep learning in particular has spawned new rodent-specific skull-stripping tools, several of which use some form of the U-Net (Ronneberger et al., 2015) as their underlying architecture. These learning-based approaches include Multi-Task U-Net (MU-Net; De Feo et al., 2021), RodentMRISkullStripping (Hsu et al., 2020), and Deep Learning-based Brain Image Processing Pipeline (DeepBrainIPP; Alam et al., 2022). MU-Net and RodentMRISkullStripping both incorporate frameworks based on a basic U-Net. DeepBrainIPP uses a network derived from the no-new U-Net (nnU-Net; Isensee et al., 2018), which is a widely-used general-purpose biomedical segmentation method.

Although deep-learning models are powerful tools for performing segmentation tasks, many of the existing deep-learning approaches require resampling of the input and output images to meet their image resolution and size requirements. Downsampling and upsampling of an image can lead to segmentation errors resulting from interpolation artifacts. DeepBrainIPP is one example of a learning-based algorithm that resamples its data to specific resolutions, which are selected based on whether the input MRI was acquired *in vivo* or *ex vivo*. In the case of *in vivo* data, DeepBrainIPP interpolates input images to 60 µm *×* 60 µm in-plane resolution with a 480 µm interplane distance. This would require an input image with 100 µm isotropic resolution, a voxel size that has been used in many studies (e.g., Bock et al., 2005; Y. Ma et al., 2008; Meyer et al., 2017), to be upsampled 1.7-fold in-plane and downsampled 4.8-fold interplane. Loss of small anatomical features in the MRI can occur with drastic downsampling, and errors propagated from the interpolation steps can reappear when the extracted brain mask is resampled back to the original MRI resolution. These methods can be made less sensitive to differences in scale without the need for resampling by including images with a variety of resolutions in the training data. This can be achieved either by expanding the dataset with images acquired at different scales or by augmenting the training data with multiple scaled versions of the original images, a technique known as scale-jittering. Still, models that use convolutional layers, which are ubiquitous in image segmentation networks (e.g., convolutional neural networks [CNNs] and U-Nets) remain sensitive to scaled patterns (Gong et al., 2014; Xu et al., 2014).

More recently, the Transformer model (Vaswani et al., 2017), which was originally developed for natural language processing (NLP), has been adapted for use in image analysis. Whereas CNNs excel at learning localized patterns, Transformers are better able to learn global context, making them less sensitive to images of different scales than convolution-based approaches are. Transformers are often built with an encoder-decoder architecture, in which the encoder learns the relationship between tokens (e.g., words in NLP or image patches in image analysis) and the decoder generates sequences of desired outputs based on the encoded inputs. One important derivative of the Transformer is the Vision Transformer (ViT) (Dosovitskiy et al., 2020). In contrast to CNNs and U-Net-based models, the ViT uses Transformer blocks rather than convolutional layers. A key element of the Transformer is its multi-head self-attention (MSA) mechanism, which allows the model to contextualize each small patch of area in the image. Conceptually, MSA helps direct the model’s attention to specific regions of the image, thereby reducing its attention to less important regions. The Transformer does this by partitioning the input image into small, non-overlapping patches (e.g., 2 pixels *×* 2 pixels) and converting these patches into embedding vectors, which are also called tokens. It then learns the sequential associations of adjacent and non-adjacent tokens.

Images can vary greatly in scale and resolution, which differs from the data addressed by natural language processing (NLP). The scale of the tokens in an NLP Transformer is fixed because each word is represented by a single token regardless of the length of the word. In contrast, a token in a ViT is derived from a small image patch of a fixed size. As a result, the tokens capture different scales when a dataset has multiple resolutions, which can make it more difficult for the model to learn a task effectively. Another challenge in using Transformers for image processing is that the task of calculating contextual information for each small image patch is computationally expensive. This issue becomes more pronounced when datasets contain large, high-resolution images. Z. Liu et al. (2021) addressed this with the Shifted Window (Swin) Transformer, which improved upon ViT by implementing a novel shifted-windowing scheme and a token merging method, which work together to compute cross-window associations efficiently and to generate a hierarchical representation of image features. This implementation enables the Swin Transformer (SwinT) to harness the benefits of the MSA mechanism of NLP Transformers and the hierarchical feature extraction capability of CNNs. These methods have been further extended for semantic segmentation by replacing the decoding scheme of Transformers with the decoding schemes of U-Nets (Ronneberger et al., 2015). Hatamizadeh et al. (2022) showed enhanced segmentation performance by using Transformer blocks as encoders and U-Net-based convolutional layers as decoders. They attributed the improvement to the incorporation of CNNs and hypothesized that convolutional layers can be more effective at evaluating local information compared to the Transformer decoders. Examples of this type of architecture include the U-Net Transformer (UNETR; Hatamizadeh et al., 2022) and the Shifted window UNETR (SwinUNETR; Hatamizadeh et al., 2021), both of which use convolutional layers in the decoding process. With the reintroduction of convolutional layers, however, the robustness to scale variance is somewhat decreased in SwinUNETR compared to the Swin Transformer.

In the present work, we developed a mouse-specific skull-stripping approach that processes images at their native resolutions. Our proposed Mouse Brain Extractor extends the SwinUNETR architecture with the goal of enhancing its robustness to scale-variance. We supplement the SwinUNETR model with additional spatial information in the form of absolute positions. We represent the positional information using Global Positional Encodings (GPEs), which we define using a shared coordinate frame based on the entire input image. The absolute positions provide additional contextual information and help improve delineation accuracy when tokens vary in scale. Our approach differs from that of SwinUNETR, which exclusively uses pairwise relationships of token positions, and from the conventional absolute positional encoding design, which defines position based on only a section of the image.

We trained and tested the Mouse Brain Extractor model using a heterogeneous set of 223 *in vivo* and *ex vivo* mouse MRI that we carefully curated from existing datasets. These data pose challenges because of variation in image resolution, image dimensions, acquisition systems and protocols used, as well as variations in brain structure due to differences in mouse strains, age, sex, disease models, and genotypes. We generated a set of reference brain segmentations for each of these MRI based on our existing lab protocol (MacKenzie-Graham et al., 2012; Meyer et al., 2023), which uses a segmentation approach that we developed previously for human data (Shattuck et al., 2001), followed by manual editing and postprocessing. We partitioned these data into three subgroups based on their degree of anisotropy and whether they were acquired *in vivo* or *ex vivo*. Each subgroup was further divided into three subsets corresponding to training, validation, and test data.

We applied our method to the test data and assessed the results relative to the manual delineations using the Dice similarity coefficient (Dice, 1945) and the 95th-percentile Hausdorff distance (HD95; Nováková et al., 2017). We also performed an evaluation study that compared our method’s segmentation results with those of nine existing tools. These included seven rodent-specific methods, namely: DeepBrainIPP, RodentMRISkullStripping, SHERM, 3D-PCNN, RATS, rBET, and antsBrainExtraction. We also trained two state-of-the-art general-purpose deep-learning segmentation methods, nnU-Net and the original SwinUNETR. Both methods were trained using the training datasets that we used to train our own method. We applied these nine methods to our test datasets and evaluated the results using Dice scores and HD95 measures. We then performed a series of two-tailed Welch’s t-tests to determine if differences between our method’s evaluation measures and those of the existing approaches were statistically significant.

## 2 Methods

### 2.1 Background

#### 2.1.1 Swin Transformer

Swin Transformers (Z. Liu et al., 2021) have shown impressive performance when processing images of differing scales at their native resolutions. The Swin Transformer takes an input image 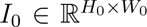, where *H*_0_ and *W*_0_ represent the height and width of *I*_0_, respectively. The image is then partitioned into non-overlapping patches of size *p×p*, where *p* is a hyperparameter. Each patch is encoded as a token and is input into a linear embedding layer, which projects the token onto an embedding space of dimension, *C*, a hyperparameter. The projected tokens then pass through multiple SwinT blocks, which are modified MSA computation blocks (Vaswani et al., 2017). The SwinT blocks compute sequential relationships among the tokens and estimate attention weights. The attention scores measure a token’s degree of influence on the other tokens in a sequence as well as its relevance in achieving the task objective (e.g., segmentation). The attention weights are adjusted during training to assign higher scores to tokens that are more influential for solving the task. The term multi-head refers to the multiple parallel self-attention calculations performed on the same input to extract multiple attention weights, which is analogous to the multiple filters used in a convolutional layer of a CNN. Self-attention is able to capture non-local information because it is computed within non-overlapping windows that are composed of *M × M* tokens. Unlike CNNs, Transformers explicitly compute the sequential relationship of the image patches in these windows, enabling the model to learn contextualized representations. After each SwinT block, the patch tokens are concatenated and the dimensionality of the resulting merged tokens is reduced using a linear layer. This decreases the number of tokens by a factor of 2, thereby creating a hierarchical structure. Cross-window associations, which enable the model to compute sequential relationships of tokens that span multiple windows, are created by shifting the windows by 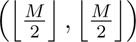 before the next set of self-attention weights are computed. As the merged tokens move further along the series of SwinT blocks, they are encoded into higher feature dimensions, which is similar to the use of increasing feature dimensions in CNNs.

These methods are implemented in neural network architectures as a repeated sequence of four steps (see Fig. 1A): a regular window-based MSA (W-MSA), a first multilayer perceptron (MLP), a shifted-window-based MSA (SW-MSA), and a second MLP. There is a residual connection before and after each step, and the output of each step is normalized using layer normalization (LN). Conceptually, W-MSA and SW-MSA determine tokens of importance, MLP provides additional network parameters and greater model flexibility, and LN and the residual connections help stabilize training. These steps are represented as a pair of Swin Transformer blocks, in which the first block computes the first two steps (W-MSA and MLP), and the second block calculates the remaining two steps (SW-MSA and MLP). In this paper, we refer to each pair of Swin Transformer blocks as a SwinT Pair. The formulation of a SwinT Pair is

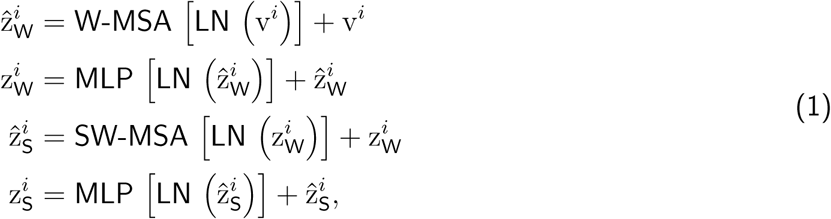

where *i* represents the index of the SwinT Pair and v*^i^* is the input to the *i*-th SwinT pair. The intermediate outputs of W-MSA and MLP for the first SwinT block are 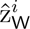 and 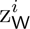, respectively. The outputs of SW-MSA and MLP in the second SwinT block are denoted by 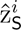 and 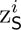. 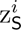 is the output for the entire SwinT Pair. The self-attention function in W-MSA and SW-MSA is defined as

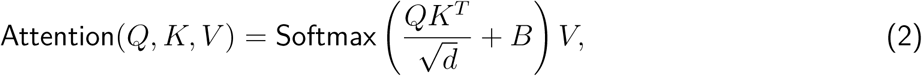

where 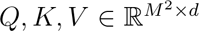 and 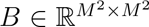 denote matrices for query, key, value, and relative position. The dimension *d* is a hyperparameter that sets the size of the key, query, value vectors, which exist as row vectors in *Q, K, V*. The scale factor 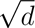 prevents small gradients during backpropagation when *QK^T^* is very large. Matrices *Q*, *K*, and *V* are derived using patch embedding vectors and are defined as

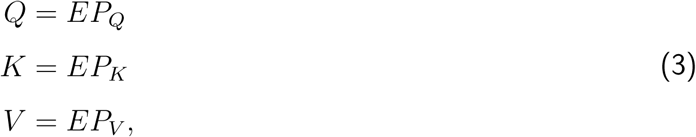

where 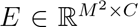 is a matrix that contains the patch embedding vectors within a window (*M × M*) and *P_Q_, P_K_, P_V_ ∈* R*^C×d^* are the network weight matrices. The Softmax in Eq. (2) estimates attention weights based on the query, key, and position matrices; these weights are then used to calculate the attention function values as weighted sums of the value matrix, *V*.

**Figure 1:**
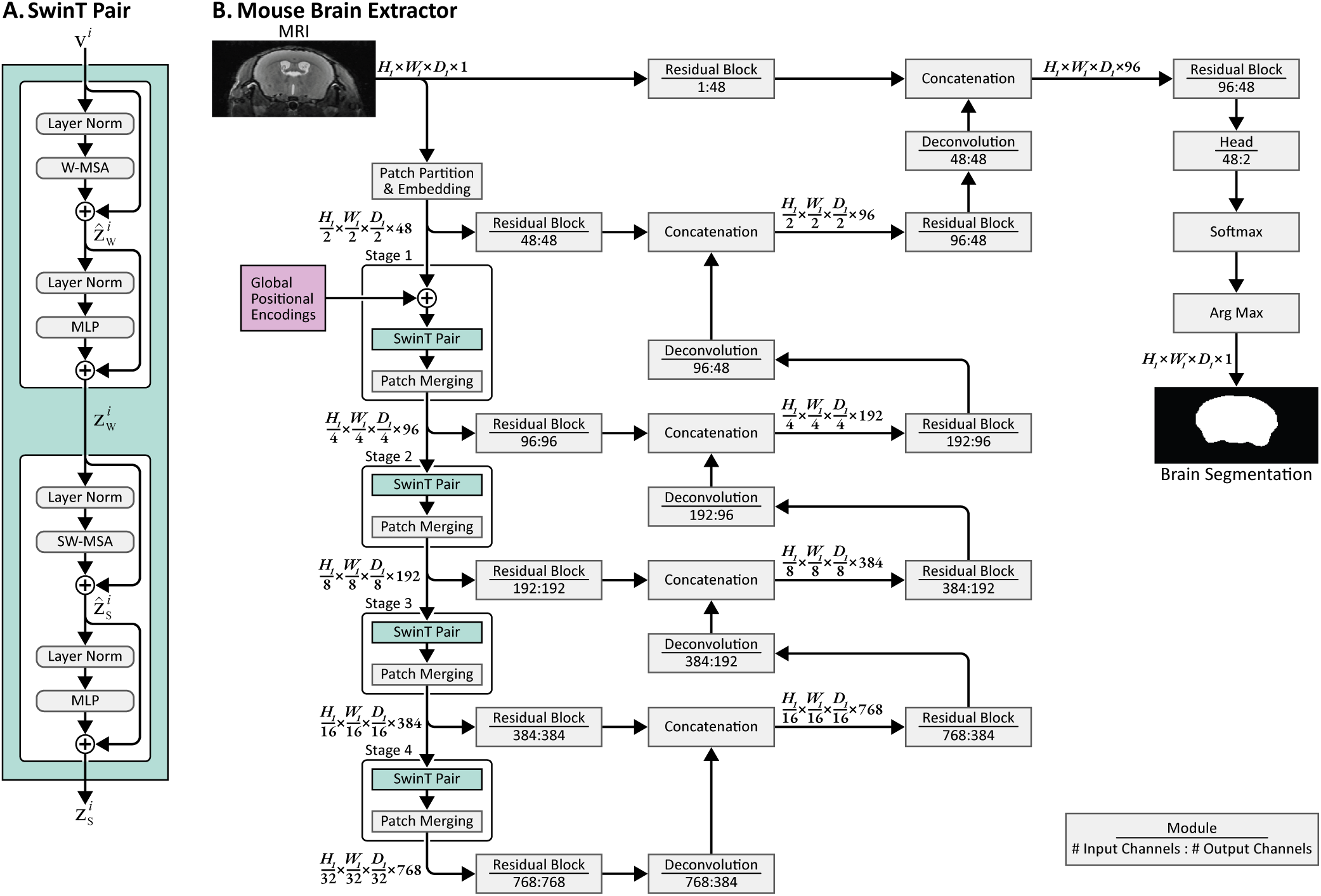
Mouse Brain Extractor Architecture. Our proposed Mouse Brain Extractor incorporates whole-image spatial information into the SwinUNETR architecture. **A. SwinT Pair:** The architectures of SwinUNETR and Mouse Brain Extractor use modules composed of two consecutive Swin Transformer blocks, which we describe as SwinT Pairs, in the encoding scheme. **B. Mouse Brain Extractor’s network architecture:** For a 3D input MRI volume (*H*_0_ *W*_0_ *D*_0_ 1), a subimage section of size *H*_1_ *W*_1_ *D*_1_ 1 is selected from the input image and partitioned into a set of 2 2 2 non-overlapping patches. These image patches are projected to an embedding space using a 1-layer CNN. Subsequently, global positional encodings are computed, downsampled by a factor of 2 in each spatial dimension, and added to the patch embedding vectors. Then, the outputs are passed into a series of SwinT Pairs, which perform similarity computations within and across windows. The encoded outputs derived at each step are input into residual and deconvolutional layers via skip connections to generate the final segmentation. For 2D models, the last dimension (*D*) is omitted. (Layer Norm: layer normalization; W-MSA: regular window multi-head self attention; MLP: multi-layer perceptron; SW-MSA: shifted window multi-head self-attention)

#### 2.1.2 SwinUNETR

The shifted window U-Net TRansformer (SwinUNETR) was developed primarily for semantic segmentation. It uses the Swin Transformer encoding scheme as its encoder and uses the upsampling deconvolutional layers from the U-Net (Ronneberger et al., 2015) as its decoder. By using convolutional layers as decoders, SwinUNETR builds upon the Swin Transformer by incorporating CNN’s ability to obtain localized information while retaining the capability of the Transformer model to capture contextualized representations. Given an image volume 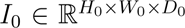, SwinUNETR samples a subimage *I*_1_ of size *H*_1_ *× W*_1_ *× D*_1_ as input to its network. We note that in some cases, one or more the dimensions of *I*_1_ may be greater than or equal to the dimensions of the input image *I*_0_. This can occur, for example, if the input image is low resolution. In such cases, the image will be zero-padded as necessary so that *I*_1_ has dimensions *I*_1_ of size *H*_1_ *× W*_1_ *× D*_1_. For simplicity, we still refer to *I*_1_ as a subimage of *I*_0_. At each iteration during training, the subimage *I*_1_ is positioned randomly within *I*_0_ to provide the model with variations of content sampled from the image. During inference, the subimage is positioned sequentially to cover the entire domain of *I*_0_.

The SwinUNETR encoder is a sequence composed of five main modules: (1) a patch partition layer; (2) a linear patch embedding layer with two consecutive Swin Transformer blocks accompanied by patch merging; and (3)-(5) three repeated modules that each consist of two consecutive Swin Transformer blocks with patch merging. The final module is followed by a bottleneck layer, which further reduces the output dimensionality while retaining relevant information. Each module that contains a SwinT Pair (i.e., modules 2-5) is defined as a stage. At each stage, the output features are passed to a CNN-decoder using skip connections, which produce the final segmentation. We note that Swin Transformers can process 2D or 3D data; for the sake of brevity, we describe 3D SwinT blocks here and in the following sections.

#### 2.1.3 Positional encoding

The patch embeddings used in Transformer models do not inherently contain spatial information, and thus the models consider them as an unordered collection of tokens. The absence of positional information can pose an issue in semantic segmentation because accurate delineation of regions often relies on the spatial relationships of image patches. In a mouse MRI, for example, image patches within the olfactory bulb may have image textures that are similar to the image textures presented by the spinal cord. Incorporating spatial information for these patches may help differentiate these anatomical regions because they are often located within distinct areas of the image.

Positional encoding was developed to add spatial information to each token explicitly. One type of positional encoding is absolute positional encoding, which encodes the absolute locations of tokens. A well-known method for absolute positional encoding was proposed by Vaswani et al. (2017) during their development of the Transformer model. They assigned a unique fixed value to each token, which encodes its absolute position using a series of sine and cosine functions with progressively increasing wavelengths. This implementation was motivated by the hypothesis that the cyclical nature of sinusoids may encourage the models to learn relative positions rather than relying on explicit numerical values (e.g., spatial Euclidean coordinates). Additionally, Vaswani et al. (2017) postulated that these sinusoidal positional encoding functions may promote the model’s generalizability towards longer inputs because they can be extrapolated more easily than learned position embeddings.

A second type of positional encoding is relative positional encoding, which is dependent on the pairwise positions of tokens. This method is used in the Swin Transformer and in SwinUNETR in the form of relative position bias matrices, denoted by *B* in Eq. (2), with learnable position embeddings. Relative positional encoding does not rely on the absolute positions of the tokens and can thus be beneficial in datasets where image features are not consistently located at the same positions. This can be particularly helpful in natural images, which can contain objects with widely diverse positions.

### 2.2 Mouse Brain Extractor

Our goal in developing Mouse Brain Extractor was to create a method that can segment the brain accurately in a mouse MRI at its native resolution. We adapted the SwinUNETR architecture to use both relative and absolute positional encoding. Our reasoning for this was based on the observation that mouse MRI are frequently acquired such that they exhibit fairly consistent placement of anatomical features within the field of view. This differs from natural images, where there is often high variability in the features of the image background or in the positioning of objects of interest.

In our Mouse Brain Extractor, we use a combination of relative position encoding and absolute position encoding to provide local and global context when processing each image patch. Our approach differs from that of SwinUNETR, which solely uses relative positional encoding, and from conventional absolute positional encoding, which derives position embeddings based only on the subimage sampled from the larger input image. Relative positional encodings are permutation invariant, which means that they are not associated with specific image features. This is useful in cases when the locations of image features vary highly across images. When subimages do not contain enough image detail, absolute positional encoding can provide the model with additional spatial information in the form of explicit token positions. However, absolute positional encoding schemes are not as beneficial in datasets with multiple scales, because they assign identical absolute positions to each subimage regardless of its location in relation to the whole image. We thus formulated Global Positional Encoding, a version of absolute positional encoding that uses a normalized coordinate system based on the full input image rather than on an input image patch.

#### 2.2.1 Global Positional Encoding

We implemented Global Positional Encoding (GPE) to be more invariant to image scale by allowing all input images to share a similar coordinate frame regardless of image size. In the case of analyzing mouse MRI, we assume that the head of the mouse is positioned near the center of the image volume and that it occupies the majority of the image space. We note that we did consider basing GPE on registration to a standard template; however, we did not observe substantial improvements using this approach during our initial testing. We thus opted not to include an extra image registration step in our approach, which would have introduced an additional dependency and a potential source of error.

The conventional approach for absolute positional encoding assigns identical absolute positions to tokens regardless of the position of the subimage *I*_1_ within *I*_0_. These absolute positional encodings are thus relative to the subimage and are not truly absolute because *I*_1_ is sampled from a different location within *I*_0_ at each iteration. Identical absolute positions could be assigned to tokens that have dissimilar image features, such as in the case of images that vary in scale.

Our approach differs in that we generate the global positional encoding for each voxel based on its location within the entire input volume, rather than just the subimage. We did this for two reasons. First, in cases where the subimage size is much smaller than the image size, e.g., for a large image with high resolution, positional information regarding the location of the subimage with respect to the whole image can provide additional information that cannot be deduced from local image patterns. This can be especially helpful if there is insufficient image detail due to reduced image contrast or low spatial frequency within the subimage. For example, a subimage of the midbrain in a low-resolution image may appear similar to a subimage of the neck muscle tissue in a high-resolution image. In scenarios similar to this, GPE may aid in correctly identifying the image patches that should be labeled as brain. Secondly, if the subimage size is larger than the size of the input image, e.g., for a small image with low resolution, the input data will be padded prior to training or inference. We do not compute GPEs in the padded areas, which is consistent with our assumption that the fields of view for whole-head mouse MRIs generally cover similar anatomical areas. This can simplify training by letting the model learn that these areas are unimportant.

We represent the GPEs as fixed 3D sinusoidal functions, which we adopted based on methods proposed by Vaswani et al. (2017), Wang and Liu (2019), and the multidim-positional-encoding software package (Tatkowski, 2024) for 1D, 2D, and 3D, respectively. This formulation produces a unique value at each token position while maintaining an identifiable pattern across the positions. We construct a GPE array, 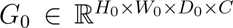, such that each voxel position defined by coordinates *x*,*y*,*z* is represented by a unique vector of length *C* = 3*c*, indexed by *k*:

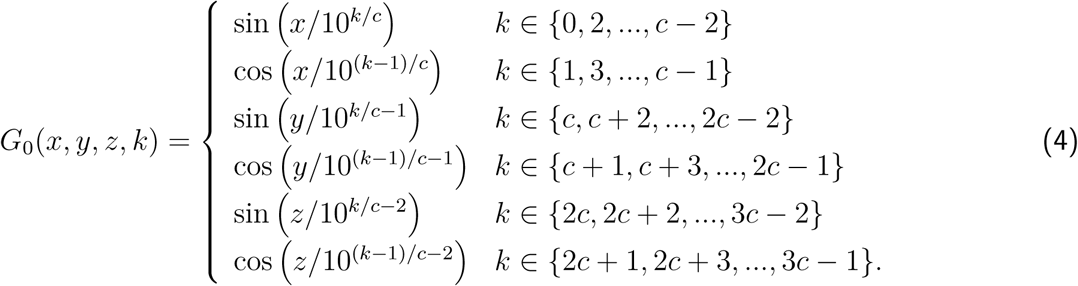

We define *C* to be a multiple of 6 so that we can partition the embedding vector into three equal segments, each of which contains values that alternate between a sine function and a cosine function. These three segments correspond to the image coordinates *x*, *y*, and *z*, which are formed by normalizing the voxel indices to the range [0,10] based on the initial input image size. This represents a set of normalized Euclidean coordinates of *I*. We empirically selected the bounds to limit the domain of the sine and cosine functions to the unit interval in radians.

Three characteristics of the GPE contribute to the generation of a distinct embedding vector at each position *x, y, z*. First, alternating sine and cosine values occur at even and odd values of *k*, respectively. Second, the period of each sinusoid increases with each increment of *k*. Third, the 3D location of the token is represented in the GPE by the three-partitioned segments.

Prior to adding the GPEs to the patch embedding vectors, we select a subregion *G*_1_ from *G*_0_, which corresponds to the region of subimage *I*_1_. We then downsample the spatial dimensions of *G*_1_ to produce 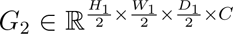 to match the spatial dimensions of the tokenized data, which are formed by reducing *I*_1_ by a factor of 2. We then select a window in *G*_2_ that corresponds to the region of the patch embedding vectors 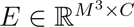 to produce 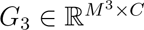. *G*_3_ is then added to *E* to generate the query, key, and value matrices, which are now defined as:

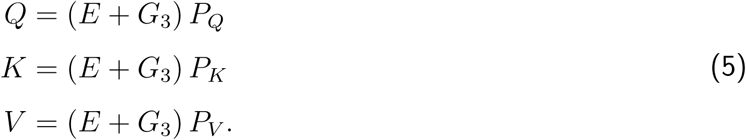

#### 2.2.2 Mouse Brain Extractor architecture and implementation

We implemented the Mouse Brain Extractor model by adapting the original SwinUNETR implementation distributed by the Medical Open Network for Artificial Intelligence Project (MONAI; Cardoso et al., 2022). The major changes that we made were the incorporation of GPEs in Stage 1 of the Swin Transformers (described in Sec. 2.2.1); the remainder of our network architecture is nearly identical to the original SwinUNETR model (Hatamizadeh et al., 2021) and differs only in input dimension and size. Figure 1 illustrates our Mouse Brain Extractor architecture, which we describe here for a 3D MRI volume input. The architecture for 2D input images is similar, but the third dimension (*D*) is omitted.

During training, we repeat the following process for each input MRI, which we represent as *I*_0_. First, the normalized Euclidean coordinates of *I*_0_ are generated and used to compute the *C*-element GPE vectors that form 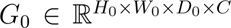 in Eq. (4), where *C* = 48. We then select a random voxel position in *I*_0_ and extract a subimage 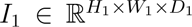 centered on this point. The subimage is formed by zero-padding or cropping each dimension if necessary to match the new size. Next, *I*_1_ is passed into a 1-layer CNN (kernel size = 2; stride size = 2; output features size = *C*) to produce 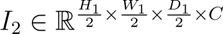 *G*_1_ is then downsampled by a factor of 2 along the spatial axes using trilinear interpolation to match the dimensions of *I*_2_, resulting in *G*_2_. We note that we only downsample in the spatial dimensions and we do not resample the *k* dimension. We also interpolate values per channel (i.e., along each *k*). This is important in maintaining the sinusoidal form in each channel. Next, *G*_2_ is added to *I*_2_ to provide absolute positions of each token. Mouse Brain Extractor, like the original SwinUNETR, has 4 stages that each contain two Swin Transformer blocks, as formulated in Eq. (1), which are followed by patch merging. Each SwinT block performs a sequence of layer normalization (LN), W-MSA, a second LN, and 2-layer MLP with Gaussian Error Linear Units (GELUs). A residual connection is added after each W-MSA and MLP. The second SwinT block is identical to the first, except it replaces W-MSA with a SW-MSA.

The patch merging process reduces the number of features by a factor of 2 and also concatenates 2*×*2*×*2 neighboring patches to form feature embeddings of size 4*C*, which are then passed into a linear layer to reduce the feature dimension to 2*C*. Thus, at each step in the encoding scheme, we have the following image resolutions: 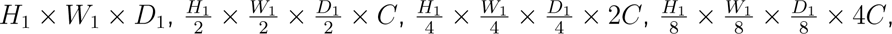 and 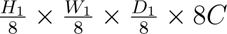

The input MRI, the patch embeddings, and outputs from Stages 1 through 3 are passed into residual blocks (2-layer CNN with kernel = 3, stride = 1, pad = 1). The resulting outputs are concatenated to the feature outputs from the previous stage, which are then deconvolved using an upsampling kernel size of 2. These concatenated outputs are passed into the residual blocks once more (2-layer CNN with kernel = 3, stride = 1, pad = 1). Stage 4 output features are input into only one set of residual blocks and passed into the deconvolutional layers. The feature dimension is reduced by half in the deconvolutional blocks. However, the deconvolutional block that restores the original resolution does not alter the feature dimension size. Lastly, the output from the decoder is passed through a single convolutional layer (kernel size = 1; stride = 1) and a Softmax function to produce the final segmentation volume (*H*_1_*×W*_1_*×D*_1_*×*2).

### 2.3 Source Data

We assembled a heterogeneous collection of *in vivo* and *ex vivo* T1- and T2-weighted MRI of mouse brains (N=223). Our motivation in creating this dataset was to be able to improve the generalizability of our method by including data that employed a variety of mouse brain anatomy and MRI protocols. In total, these data were collected using six types of scanners, six different T2-weighted pulse sequences, and two different T1-weighted pulse sequences. Our dataset comprised 163 *in vivo* and 60 *ex vivo* mouse images. These included healthy brains (N=69), brain models of diseases and disorders (N=106), and physically injured brains (N=48). We gathered this dataset from images that we acquired previously in separate studies (Hubbard et al., 2021; Itoh et al., 2023; MacKenzie-Graham et al., 2009; Meyer et al., 2017), that were acquired as part of ongoing studies of specific mouse models (Tabbaa et al., 2023), or that we retrieved from an open-access repository (Y. Ma et al., 2008). All studies from which we retrieved data were performed in accordance with the National Institutes of Health Guide for the Care and Use of Laboratory Animals and appropriate Institutional Animal Care and Use Committees (IACUC). A summary of the imaging data and their corresponding demographic information is shown in Table 1. Additional details regarding the MR scanners, imaging sequences, image dimensions, and scanning resolutions are provided in Appendix Table 4.

**Table 1:** Dataset demographics. We selected a heterogeneous collection of mouse MRI containing a mixture of populations, strains, genotypes, ages, and sexes for training and evaluation. The data were acquired previously as part of other studies (Hubbard et al., 2021; Itoh et al., 2023; MacKenzie-Graham et al., 2009; Meyer et al., 2017), acquired as part of ongoing studies of specific mouse models (Tabbaa et al., 2023) or retrieved from an open-access repository (Y. Ma et al., 2008). (DPI: days-post-injury indicates the number of days after brain injury or sham surgery, EAE: experimental autoimmune encephalomyelitis, TBI: traumatic brain injury, WT: wild type, HET: heterozygous type, CKO: conditional gene knockout, SOD1: superoxide dismutase 1, AAV: adeno-associated virus)

| Dataset | Cohort | Strain | Genotype | Sex | Age at Scan (mos. ± sd) | Notes |
| --- | --- | --- | --- | --- | --- | --- |
| *in vivo* isotropic | EAE<br>Meyer et al., 2017 | 48 C57BL/6 | 48 WT | 25 F / 23 M | 4.94 ± 0.48 | 30 EAE / 18 healthy controls |
| | Aging<br>Itoh et al., 2023 | 26 C57BL/6 | 26 WT | 13 F / 13 M | 13.65 ± 6.86 | Aged 5 to 24 months |
| | NeAt<br>Ma et al., 2008 | 5 C57BL/6 | 5 WT | 0 F / 5 M | 3.25 ± 0.25 | |
| *in vivo* anisotropic | *Chd8*<br>Tabbaa et al., 2023 | 21 B6-CC61<br>15 B6-CC17 | 18 HET<br>18 WT | 18 F / 18 M | 10.43 ± 1.53 | |
| | TBI<br>Hubbard et al., 2021 | 48 C57BL/6 | 48 WT | 24 F / 24 M | 4.97 ± 2.17 | 24 injured / 24 sham<br>12 0-DPI / 12 30-DPI / 12 100-DPI / 12 166-DPI |
| *ex vivo* | EAE<br>MacKenzie-Graham et al., 2009 | 60 C57BL/6 | 48 WT<br>12 CKO | 47 F / 13 M | 4.67 ± 1.27 | 40 EAE / 20 healthy controls<br>12 SOD1<sup>-/-</sup><br>12 AAV-treated |

**Table 2:** Training data. Training data for nnU-Net, SwinUNETR, and Mouse Brain Extractor included uncorrected and N4-corrected MRI data. Data augmentation was used to generate images with severe artificial bias field artifacts and scale-jittering. The scale-jittered data were not included in the training dataset for the nnU-Net models because nnU-Net’s workflow resamples the data to the median resolution of the images on which it was trained (see Sec. 2.9.1). On-the-fly data augmentation was performed during training to further expand the dataset and is described in Section 2.5.1.

| Dataset | Model | Uncorrected/<br>N4-corrected | Artificial bias<br>field artifact | Scale-jittered<br>(Uncorrected/<br>N4-corrected) | Total |
| --- | --- | --- | --- | --- | --- |
| *in vivo* anisotropic | 2D nnU-Net | 56 / 56 | 8 | None | 120 |
| *in vivo* anisotropic | 3D full resolution nnU-Net | 56 / 56 | 8 | None | 120 |
| *in vivo* isotropic | 2D nnU-Net | 54 / 54 | 8 | None | 116 |
| *in vivo* isotropic | 3D full resolution nnU-Net | 54 / 54 | 8 | None | 116 |
| *ex vivo* | 2D nnU-Net | 40 / 40 | 5 | None | 85 |
| *ex vivo* | 3D cascade nnU-Net | 40 / 40 | 5 | None | 85 |
| *in vivo* anisotropic | 2D SwinUNETR | 56 / 56 | 8 | 12 / 12 | 120 |
| *in vivo* isotropic | 3D SwinUNETR | 54 / 54 | 8 | 14 / 14 | 116 |
| *ex vivo* | 3D SwinUNETR | 40 / 40 | 5 | 10 / 10 | 85 |
| *in vivo* anisotropic | Mouse Brain Extractor | 56 / 56 | 8 | 12 / 12 | 120 |
| *in vivo* isotropic | Mouse Brain Extractor | 54 / 54 | 8 | 14 / 14 | 116 |
| *ex vivo* | Mouse Brain Extractor | 40 / 40 | 5 | 10 / 10 | 85 |

We subdivided the data into three groups (see Table 1): *in vivo* isotropic, *in vivo* anisotropic, and *ex vivo*. Images in the anisotropic *in vivo* group had a voxel aspect ratio in the range [3.98, 15.36], where we define voxel aspect ratio as the maximum edge length of a voxel divided by its minimum edge length. Aspect ratios in this range are not uncommon in preclinical imaging, where *in vivo* brains are often imaged with high in-plane resolution and a corresponding larger slice thickness. The *ex vivo* images were acquired with a smaller range of voxel aspect ratios (1–1.85) and did not exhibit large partial volume effects. Because these images were mostly uniform in their appearance, we opted to have a single group for the *ex vivo* data.

While one could in principal use all three groups of data to train a single model, we opted to produce separate models for each type of data. A primary aim of the model is to correctly characterize the distribution of the input datasets. However, distinct dissimilarities in image appearance across the datasets as well as the more subtle differences within datasets result in a data distribution with many modes. The complexity of the distribution-fitting process increases with the number of modes and has the potential to decrease the trained model’s accuracy. Thus, a simpler data distribution with less data variability makes it easier to parameterize the distribution accurately and enhances the model’s performance. In our data, we have a mixture of voxel aspect ratios and *in vivo* and *ex vivo* preparations, which all contribute to variations in image appearance. While highly anisotropic images often have very high resolution in-plane, they can still exhibit blurry brain boundaries due to the partial volume effect. Each voxel is an average of the signals within its volume, and a large slice thickness will thus cause voxels to average signals from multiple tissue types. This makes it more difficult to distinguish between non-brain and brain and to localize the brain boundary accurately. Additionally, compared to *ex vivo* images, *in vivo* images typically have lower image resolution and contrast because of limited scan times. Images of *ex vivo* models can be acquired with much higher resolution and quality, and may thus capture finer anatomical details and boundaries. Moreover, *ex vivo* brains may have a distorted shape compared to *in vivo* brains because the ventricles collapse due to reduced fluid pressure.

#### 2.3.1 *In vivo* isotropic MRI

The *in vivo* isotropic data comprised scans from 79 C57BL/6 wildtype (WT) mice. These included 48 mice from a study investigating experimental autoimmune encephalomyelitis (EAE), a mouse model of multiple sclerosis (Meyer et al., 2017), 26 mice from a normative aging study (Itoh et al., 2023), and 5 scans from the open-access MR Microscopy (MRM) NeAt repository (Y. Ma et al., 2008). MRI from the EAE and aging studies were acquired at the Ahmanson-Lovelace Brain Mapping Center (ALBMC) at UCLA and at the Small Animal Imaging Core (SAIC) at Children’s Hospital Los Angeles (CHLA). The MRI data from the MRM NeAt dataset were retrieved from the DigitAl Medicine Analytic (DAMA) Lab’s GitHub repository (D. Ma, 2020) that were imaged at the Brookhaven National Laboratory and National High Magnetic Field Laboratory (Y. Ma et al., 2008).

#### 2.3.2 *In vivo* anisotropic MRI

The *in vivo* anisotropic data included 84 MRI from two separate studies: 36 from a study involving mice with a loss-of-function mutation (haploinsufficiency) in *Chd8*, a high-confidence autism spectrum disorder (ASD) risk gene (Tabbaa et al., 2023), and 48 from a multi-site traumatic brain injury (TBI) study (Hubbard et al., 2021).

The Chd8 data included images of 21 B6-CC61 and 15 B6-CC17 mice. These two strains were selected because B6-CC61 exhibits high Chd8 mutation vulnerability resulting in atypical social behavior compared to the resilient B6-CC17 strain (Tabbaa et al., 2023). Each mouse was scanned using two sequential acquisitions that produced two separate images that contained the anterior and posterior halves of the head. This acquisition method was adopted to adjust the repetition time (TR) to acquire an acceptable contrast, which also provided an acceptable resolution and total scan time. As described in Sec. 2.4, these pairs of images were later combined to form single whole-head images for each mouse brain.

The TBI data included MRI of C57BL/6 mice that received severe controlled cortical impact (CCI) injury or sham surgery prior to scanning. Both sets of mice received a craniotomy on the left side of the skull, but only the CCI group received an injury using a 3 mm flat-tip impactor at the same location. Following the procedure, a plastic surgical disc was placed on the site of the surgery. The mice were imaged at 0, 30, 100, and 166 days post CCI or sham surgery.

#### 2.3.3 *Ex vivo* MRI

All 60 *ex vivo* MRI (47F/13M; 48 WT/12 conditional knock-out [CKO]; age at scan 4.67±1.27 months) were collected from a mixture of C57BL/6 healthy and EAE-induced mice as part of a previous study on EAE (MacKenzie-Graham et al., 2009). These *ex vivo* images were scanned using T1- and T2-weighted sequences at the Duke Center for In Vivo Microscopy (CIVM) (N=12), the Beckman Institute at the California Institute of Technology (N=24), and SAIC at CHLA (N=24). They consisted of isotropic images and anisotropic images with voxel dimension ratios in the range 1.33–1.85.

The T2-weighted images acquired at CIVM were collected from a female cohort (age at scan 4.50±1.40 months) and included four EAE-induced mice and eight healthy controls with a range of 63 to 147 days post-EAE induction. Twelve mice (N=12; 12F; 5 EAE/7 healthy controls; age at scan 4.25±0.70 months; post-EAE induction range of 15 to 55 days) scanned at the Beckman Institute were imaged using a T2-weighted imaging protocol, while the other twelve (N=12; 12F; 7 EAE/5 healthy controls; age at scan 3.40±0.69 months; post-EAE induction range of 20 to 80 days) were acquired using a T1-weighted imaging sequence.

The *ex vivo* cohort scanned at SAIC consisted of 12 transgenic mice with a conditional knockout of the superoxide dismutase (SOD1) enzyme-encoding gene (SOD1^-/-^) (7F/5M; 12 EAE; age at scan 6.48±0.33 months; 41 days post-EAE induction) and 12 mice treated with adeno-associated virus (AAV) (4F/8M; 12 EAE; age at scan 4.7 months; 43 days post-EAE induction).

### 2.4 Data preparation

The MRI data for this study were selected to capture variability in head and neck shape and position, brain size due to strain and genotype, field inhomogeneity, ringing artifacts, RF interference, and anatomical abnormalities. After selecting an initial set from each study through visual inspection, we identified additional images of similar image intensity distributions by computing the Jensen-Shannon distances between the selected images and the potential candidate images. We identified subsets of these images to be used as training, validation, and test datasets.

#### 2.4.1 MRI volume preprocessing

Each MRI was reoriented such that its image data were stored in a consistent right-anterior-superior orientation. We note that this was achieved by permuting the axes to reorder the data, which does not require resampling or interpolation. We then performed limited pre-processing on the training, validation, and test datasets. For most datasets, this involved only bias field correction. The *Chd8* data required processing to combine the two acquisition frames, which we achieved as follows. We first concatenated the two sections to create a single image. However, differing bias field effects produced a distinct discontinuity where the images were joined. We addressed this by performing slice-based normalization to create a smoother transition between the two sections. The normalization was performed by: (1) computing the average intensity value per slice 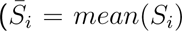, where *S_i_* denotes the intensity values in image slice index *i*); (2) fitting a 2nd degree polynomial model on *S̅* to obtain estimated intensity values *Ŝ* using the function’s coefficients; and (3) modifying the intensity values by 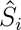 per slice 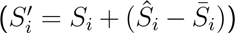. The resulting concatenated image was treated as one MRI volume in subsequent processing and analysis. We then performed bias field correction on all image volumes using the N4 method from ANTs (Tustison et al., 2010). We refer to these data as the N4-corrected images in this manuscript, while we refer to the original set as uncorrected.

#### 2.4.2 Delineation

We created a reference brain mask for each MRI volume following in-house protocols developed by A.M.G. (MacKenzie-Graham et al., 2012; Meyer et al., 2023). We generated an initial set of brain masks for all MRI volumes using BrainSuite’s Brain Surface Extractor (BSE; version 21a), a skull-stripping tool that we originally developed for T1-weighted MRI of the human brain (Shattuck et al., 2001). We applied BSE to the N4-corrected images and used its automated parameter selection feature, which optimizes algorithm settings using a cost function designed for human brains (Rajagopal et al., 2017). When needed, we manually adjusted the program settings and reapplied BSE to generate improved brain masks. A trained rater (H.H.) with 4 years of brain labeling experience then overlaid the masks on the N4-corrected images and performed manual edits using the mask editing tool in the BrainSuite GUI (Shattuck & Leahy, 2002). The brain masks included all brain tissue and excluded the trigeminal nerve, CSF ventral to the pons, and the optic nerve. Any spinal cord tissue present posterior to the last coronal slice where the cerebellum is not present was also removed from the mask. Manual corrections required approximately 10 mins per MRI for the *in vivo* anisotropic data, 30 mins per MRI for the *in vivo* isotropic data, and 60 mins per MRI for the *ex vivo* data.

While manual edits can represent a gold standard for comparison, they are prone to some errors and irregularities. In this work, the completed masks exhibited uneven boundaries across slices because the brain masks were edited on a slice-by-slice basis primarily in the coronal plane. There were some additional errors produced by stray mouse clicks, which resulted either in holes interior to the mask or extraneous clusters of voxels that were disconnected from the brain mask.

We addressed these issues by applying a series of processing steps to all mask volumes in the training, validation, and test datasets. We corrected for stray mouse clicks by selecting the largest connected component in the image as the brain mask and then filling any holes in that mask. We smoothed uneven segmentation boundaries across slices by first smoothing the mask with a 3D Gaussian filter with *σ*=1 voxel and then thresholding the smoothed image at 0.5 to produce a binary mask. An expert rater (A.M.G.) reviewed the final brain masks visually and approved them.

Examples of the final manually-edited masks for *in vivo* isotropic, *in vivo* anisotropic, and *ex vivo* data are shown in Fig. 2. Distinctive characteristics for each type of image can be observed, some of which may present difficulties when determining boundaries. In the *in vivo* isotropic data, lower contrast and attenuated signals in the ventral region resulting from field inhomogeneity make brain edge detection challenging for both human raters and automated methods. Although N4 correction partially restores the signals and contrast, this area often remains hypointense and difficult to delineate. Another problematic feature can be observed in the *in vivo* anisotropic data, where the partial volume effect from the larger slice thickness blurs the brain boundaries. This is particularly visible in the rostral regions where the shape of the brain changes more dramatically. These issues are less commonly observed in *ex vivo* data, which typically have higher resolutions and less field inhomogeneity. However, in the case of some of the T1-weighted *ex vivo* images, brain boundaries are less distinct and still pose challenges.

**Figure 2:**
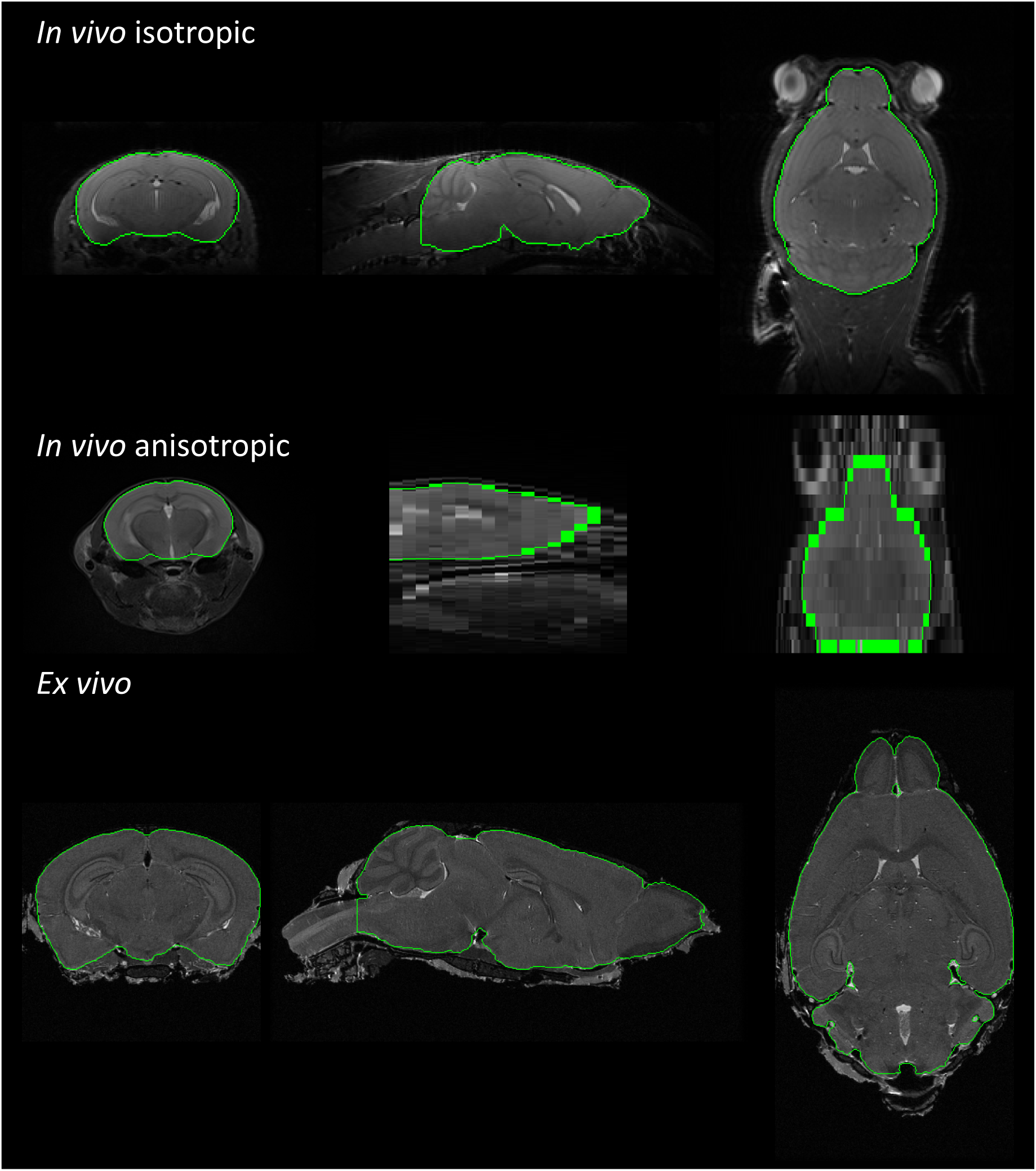
Manually-edited delineations with postprocessing. An example of an uncorrected image from each dataset is shown in each row. Top: *in vivo* isotropic image of an EAE-induced female from the EAE cohort; middle: *in vivo* anisotropic image of a sham-surgery female from the TBI cohort; and bottom: *ex vivo* image of an EAE-induced female from the EAE cohort. Green contours indicate the brain delineation boundaries, which were initially generated with BSE (Shattuck et al., 2001), corrected manually according to the protocol described by MacKenzie-Graham et al. (2012) and Meyer et al. (2023), and minimally postprocessed (see Sec. 2.4.2). Some of the characteristics that pose challenges for automated segmentation include low contrast due to field inhomogeneity (*in vivo* isotropic) and blurring of anatomical boundaries due to larger slice thicknesses (*in vivo* anisotropic). Higher resolution images (*ex vivo*) may provide clearer details that improve automated segmentation results.

### 2.5 Training, Validation, and Testing Datasets

We partitioned each of the three groups of data in Table 1 (*in vivo* isotropic, *in vivo* anisotropic, and *ex vivo*) into three subsets for training, validation, and testing. The training and test datasets were organized to exhibit similar heterogeneity in demographics and image appearance. We made use of both the uncorrected and N4-corrected data in our training and evaluation. The delineated mask for each MRI volume was used as the ground truth segmentation for both the uncorrected and N4-corrected images. We included the N4-corrected images because the N4 method is frequently performed as a preliminary step in image processing pipelines and users may prefer to run brain extraction on the corrected data. We also used the N4-corrected data to represent images with minimal bias field artifacts. We note that the N4 method is not guaranteed to produce images completely free from bias field effects, thus the N4-corrected images exhibit a range of small field inhomogeneity artifacts that help improve the generalizability of our model.

#### 2.5.1 Training Data

For the *in vivo* isotropic dataset, MRI from 56 mice were used for training: 33 from the *in vivo* EAE dataset (33 C57BL/6; 33 WT; 17F/16M; age at scan 4.94±0.48 months; 12 EAE (45 days post EAE-induction)/21 healthy controls; 20 from the *in vivo* aging dataset (20 C57BL/6; 20 WT; 10F/10M; age at scan 13.80±6.72 months); 3 from MRM NeAt (3M; 3 C57BL/6). For the *in vivo* anisotropic dataset, MRI from a total of 56 mice were used: 24 from the *Chd8* cohort (14 B6-CC61/10 B6-CC17; 12 HET/12 WT; 12F/12M; age at scan 10.30±1.55 months); and 32 from the TBI cohort (32 C57BL/6; 32 WT; 16F/16M; age at scan 4.97±2.18 months; 16 injured/16 sham surgery; 8 from each of 0, 30, 100 and 166 days post injury. The *ex vivo* dataset used data from 40 mice from the *ex vivo* EAE dataset (40 C57BL/6; 32 WT/8 CKO; 30F/10M; age at scan 4.67±1.25 months; 13 EAE [30.22±24.40 days post EAE-induction]/27 healthy controls; 8 SOD1^-/-^; 8 AAV-treated). The N4-corrected images of each of these MRI were added to the training dataset.

##### Data augmentation

We included pre-computed data augmentations in the training datasets, and we also performed on-the-fly data augmentation during training. The pre-computed augmentations consisted of data with artificial bias fields and resolutions not present in the original training data. We chose to precompute these particular operations because they are very compute-intensive, as compared to the augmentations that we did compute on the fly. By precomputing them, we generate them only once and reduce computational load during training. The on-the-fly data augmentations were generated randomly at each iteration during training, which greatly increased the effective size of the training dataset. Details regarding the on-the-fly data augmentation methods are provided in Section 2.6.

We performed two data augmentation transforms on two disjoint subsets of training data prior to training. On the first subset, we applied bias field transforms which we represented as smoothly-varying 4th-degree polynomial functions with coefficients ranging from 0.2 to 0.4. We applied these fields to 8 *in vivo* isotropic, 8 *in vivo* anisotropic, and 5 *ex vivo* images.

For the second subset, we resampled the MRI to 12 distinct resolutions to enhance Mouse Brain Extractor’s ability to to learn the associations between GPEs and tokens. We hypothesized that by supplementing the training data with data containing a variety of resolutions not present in the original data, Mouse Brain Extractor can learn the relationship between GPEs and image patches. A subset of MRI volumes from the training data were resampled to 12 different resolutions using cubic spline interpolation. The corresponding mask volumes were resampled using a technique similar to nnU-Net’s interpolation method and consisted of converting the binary labels into one-hot encoding vector images, performing linear interpolation across the channels, and taking the argmax (index of the maximum value) of each vector (Isensee et al., 2018). For the *in vivo* isotropic resolution datasets, a subset of 28 images from the *in vivo* EAE, aging, and MRM NeAt datasets (14 native/14 N4-corrected) were resampled to 50 µm, 75 µm, 150 µm, and 200 µm isotropic resolution. For the *in vivo* anisotropic resolution data, a subset 24 images from the *Chd8* and TBI datasets (12 native/12 N4-corrected) were resampled in-plane to 60 µm, 100 µm, 150 µm, and 200 µm isotropic resolutions in-plane (coronal plane). For the *ex vivo* data, a subset of 20 images from the *ex vivo* EAE dataset (10 native/10 N4-corrected) were resampled to 20 µm, 25 µm, 30 µm, and 50 µm isotropic resolution.

#### 2.5.2 Validation data

We created our validation dataset by selecting at random one-fifth of the combined set, which included both the training data described in 2.5.1 and the precomputed data augmentations. This subset was used to assess the generalizability of our model during training and to determine post-processing methods. The resulting validation dataset consisted of 10 MRI for the *in vivo* isotropic dataset, 10 MRI for the *in vivo* anisotropic dataset, and 5 MRI for the *ex vivo* dataset.

#### 2.5.3 Test data

The remaining data that were not included in the training or validation datasets were used as the test dataset. A summary of the demographic information of the test datasets are outlined in Appendix Table 5. For the *in vivo* isotropic resolution datasets, 15 MRI were drawn from the *in vivo* EAE dataset (15 C57BL/6; 15 WT; 8F/7M; age at scan 4.91±0.48 months; 6 EAE (45 days post EAE-induction)/9 healthy controls), 6 were drawn from the aging dataset (6 C57BL/6; 6 WT; 3F/3M; age at scan 13.17±7.94 months), and 2 MRI were drawn from MRM NeAt dataset (2M; 2 C57BL/6). For the *in vivo* anisotropic resolution datasets, a total of 28 MRI were drawn: 12 from the *Chd8* dataset (7 B6-CC61/5 B6-CC17; 6 HET/6 WT; 6F/6M; age at scan 10.68±1.52 months) and 16 from TBI dataset (16 C57BL/6; 16 WT; 8F/8M; age at scan 4.97±2.21 months; 8 injured/8 sham surgery; 4 from each of 0, 30, 100 and 166 days post injury). The voxel aspect ratio ranges were identical to the ones in the training datasets and varied between 3.98 and 15.36. From the *ex vivo* datasets, all 20 images were taken from the *ex vivo* EAE dataset (20 C57BL/6; 16 WT/4 CKO; 17F/3M; age at scan 4.66±1.34 months; 13 EAE (51.38±29.90 days post EAE-induction)/7 healthy controls; 4 SOD1^-/-^; 4 AAV-treated). Four of these images had a voxel aspect ratio of 1.33, while the remaining 16 had isotropic resolutions.

### 2.6 Training

We trained three separate Mouse Brain Extractor models, which were each dedicated to a particular type of data: *in vivo* isotropic, *in vivo* anisotropic, and *ex vivo*. For the *in vivo* anisotropic model, we used an input subimage size of 128 *×* 128 (*H*_1_ = *W*_1_ = 128) in the coronal view because the coronal plane had the highest resolution in this dataset. During development, we observed better performance with a 2D model than with a 3D model, the accuracy of which may have been reduced by the low resolution in the sagittal and axial planes. For the *in vivo* isotropic and *ex vivo* datasets, we used an input subimage size of 96 *×* 96 *×* 96 (*H*_1_ = *W*_1_ = *D*_1_ = 96), because 3D models showed better performance than 2D models. In these 3D models, we selected 96 voxels instead of 128 to reduce GPU memory usage so that the models could be run within the 24GB available on our systems.

The maximum iterations used for each model were 90,000 for *in vivo* isotropic models, 55,000 for *in vivo* anisotropic models, and 90,000 for *ex vivo* resolution models. We used an initial learning rate of 5 × 10^−4^ with a linear warm-up with a cosine annealing learning rate scheduler (Loshchilov & Hutter, 2016). The models with the best Dice scores on the validation datasets were chosen as the final models. During training, additional data transformations were randomly performed before each iteration for all models. The transforms were similar to those used by nnU-Net Version 2 (Isensee, 2024) and included: size scaling (multiplicative factor in [0.7,1.4]); rotations ([-90°,90°]); gamma scaling (with and without inversion; *γ* = [0.7,1.5]); flipping across all orthogonal axes; Gaussian noise (*σ*^2^ *∈* [0,0.1]); Gaussian smoothing (*σ ∈* [0.5,1]); intensity scaling (multiplicative factor in [0.75,1.25]); contrast augmentation (contrast range multiplicative factor in [0.75,1.25]); simulation of low resolution (zoom factor in [0.5,1]); downsampling using nearest-neighbor interpolation and upsampling using cubic spline interpolation); and translations ([-50, 50] voxels on all axes). The probabilities for each transformation varied and were independent of each other.

### 2.7 Inference and Post-processing Steps

During inference, we used a tile overlap strategy, which runs inference on a section of image at a time, with the section being the size of the subimage used by the network. Each subsequent section overlaps the previous section by 0.8. The final segmentation is determined by the majority count.

The last stage of our segmentation approach is a set of post-processing steps designed to improve the outputs from Mouse Brain Extractor. During the development of our method, we reviewed results for the validation datasets and observed some minor errors in a small subset of the predicted labels. These included small holes internal to the brain mask, disconnected mask segments, and rough boundaries. We addressed these by developing a post-processing sequence that selects the largest connected component and fills any cavities internal to it. It then smooths the brain boundaries by applying a Gaussian filter (*σ* = 1 voxel) and then thresholding at 0.5 to yield a binary mask. This smoothing step was applied to the *in vivo* anisotropic and *ex vivo* model outputs. We did not apply Gaussian smoothing to the *in vivo* isotropic model outputs because it produced worse Dice scores on the validation data.

### 2.8 Evaluation

We evaluated our method’s segmentation performance using the test data described in Section 2.5.3. We compared each output with the corresponding manually-edited segmentation using Dice similarity (Dice, 1945) and 95th-percentile Hausdorff distance (HD95; Nováková et al., 2017) measures. Dice scores range from 0 to 1, with higher scores indicating a greater overlap between the automated method and manual segmentation (1 indicates perfect agreement; 0 indicates no agreement). The symmetric Hausdorff distance (HD), for a given segmentation *T* and *U*, is the maximum of the minimum distances from the boundary voxels of *T* to *U* and from the boundary voxels of *U* to *T*. Smaller Hausdorff distances indicate closer alignment of the segmentation boundaries. Here, we use the HD95 measure, which is the value at the 95th percentile of the minimum distances. The usage of the 95th percentile instead of the maximum distance makes the measure more robust to the effect of outliers.

### 2.9 Comparison with Existing Approaches

We performed an evaluation comparing our method’s segmentation results with those of nine existing approaches. The methods we compared were nnU-Net (Isensee et al., 2018), SwinUNETR (Hatamizadeh et al., 2021), DeepBrainIPP (Alam et al., 2022), RodentMRISkullStripping (Hsu et al., 2020), SHERM (Y. Liu et al., 2020), 3D-PCNN (Chou et al., 2011), RATS (Oguz et al., 2014), rBET (Wood et al., 2013), and antsBrainExtraction (Avants et al., 2011). We assessed their performance by computing Dice and HD95 measures of their segmentation outputs relative to the manually-edited masks. We then performed statistical comparisons of these measures against our own results.

#### 2.9.1 Processing with Existing Approaches

For each method, we processed the test images after preparing them based on the pre-processing steps reported in each method’s corresponding publication or usage instructions. We describe the individual preparations used for each method in more detail below. In a subset of cases, we performed additional steps (e.g., cropping of images or modification of method parameters) to meet method-specific usage requirements or to produce satisfactory segmentations. Table 3 summarizes the methods that were compared and the datasets on which they were evaluated.

**Table 3:** Evaluation Summary. Shown are the methods that were compared and the datasets on which they were evaluated. Some methods were not evaluated on all data subsets because of their input requirements (see Sec. 2.9.1). Incomplete results were obtained for 3D-PCNN, which terminated with an error on four images from the N4-corrected *in vivo* anisotropic dataset, and for DeepBrainIPP, which produced empty brain masks for six uncorrected and five N4-corrected images from the *in vivo* anisotropic datasets.

|  | Uncorrected |  |  | N4-corrected |  |  |
| --- | --- | --- | --- | --- | --- | --- |
|  | In vivo iso. | In vivo aniso. | Ex vivo | In vivo iso. | In vivo aniso. | Ex vivo |
| SHERM |  |  |  | Evaluated | Evaluated |  |
| RodentMRISkullStripping | Evaluated | Evaluated |  | Evaluated | Evaluated |  |
| 3D-PCNN |  |  |  | Evaluated | Incomplete Results |  |
| RATS |  |  |  | Evaluated | Evaluated |  |
| antsBrainExtraction | Evaluated | Evaluated | Evaluated |  |  |  |
| DeepBrainIPP | Evaluated | Incomplete Results | Evaluated | Evaluated | Incomplete Results | Evaluated |
| rBET | Evaluated | Evaluated | Evaluated | Evaluated | Evaluated | Evaluated |
| nnU-Net | Evaluated | Evaluated | Evaluated | Evaluated | Evaluated | Evaluated |
| SwinUNETR | Evaluated | Evaluated | Evaluated | Evaluated | Evaluated | Evaluated |
| Mouse Brain Extractor | Evaluated | Evaluated | Evaluated | Evaluated | Evaluated | Evaluated |

Evaluated Incomplete Results Not Evaluated

##### nnU-Net

We trained nnU-Net using the same training dataset that we used for Mouse Brain Extractor, with the exception of the precomputed augmented data that were produced by scale-jittering. nnU-Net automatically resamples its input data to the median resolution of the datasets on which it is trained (Isensee et al., 2018), which means that these augmented images would be resampled back to their original resolutions. This is similar to nnU-Net’s built-in low-resolution simulation data augmentation, thus we deemed it unnecessary for nnU-Net’s training. The presence of many additional resampled images in the training dataset could possibly prevent the nnU-Net models from optimizing on the original, non-resampled images.

All nnU-Net models were trained using their default method configurations and settings, apart from small changes in the data augmentation methods. We modified the degrees of image rotation used in nnU-Net’s data augmentation to match those of our proposed models. Additionally, we modified the probabilities that determine the likelihood of a specific transformation being applied to more closely match natural occurrences. For example, left-right (LR) flips are more frequently observed than up-down (UD) flips in MR data, so we adjusted the LR mirroring probability to be higher than that of UD mirroring.

NnU-Net outputs training recommendations based on the training dataset’s image properties, such as image sizes and resolutions. Based on these instructions, we employed a 5-fold validation technique and trained the 2D U-Net and 3D full-resolution U-Net for all datasets. For *ex vivo* data, we trained an additional configuration, the 3D cascade U-Net, which was recommended by nnU-Net due to the high image resolution and size. Each model was trained for 1000 iterations. After assessing the performance of each model configuration using the validation data, nnU-Net selected the 2D U-Net as the best configuration for all dataset types (Isensee et al., 2018). NnU-Net also determined post-processing methods for each model: none for *in vivo* anisotropic, and selection of the largest connected component for *in vivo* isotropic and *ex vivo* models.

##### SwinUNETR

We trained SwinUNETR identically to how we trained Mouse Brain Extractor. Specifically, the training and validation datasets, data augmentations, training hyperparameters (e.g., number of iterations and learning rates), optimizers, learning rate schedulers, and validation methods were the same as we used for Mouse Brain Extractor. During inference, we also used a tile overlap strategy with an overlap ratio of 0.8 and the same post-processing that we described in Sec. 2.7.

##### DeepBrainIPP

DeepBrainIPP (Alam et al., 2022) provides multiple pre-trained weights. We selected its *invivo-2* and *exvivo-1* models, which most closely matched the resolutions and types of our datasets. Because their models were trained with uncorrected and N4-corrected images, we included both sets in DeepBrainIPP’s evaluation. However, due to input image size restrictions, we manually cropped the images so that the pipeline could be run. We also performed extra steps external to DeepBrainIPP to prepare the segmentation outputs for evaluation. As part of its workflow, DeepBrainIPP performs skull stripping and paraflocculi segmentation separately and outputs two individual masks for these two regions. We thus combined the two segmentation masks into a single binary mask (see App. B for details).

##### RodentMRISkullStripping

We applied RodentMRISkullStripping (Hsu et al., 2020) to both the original and the N4-corrected *in vivo* isotropic and anisotropic test datasets. We excluded *ex vivo* datasets because the developers of this method only trained it on *in vivo* data.

##### SHERM

We applied SHERM (Y. Liu et al., 2020) to all of the N4-corrected *in vivo* data. We excluded the *ex vivo* datasets because SHERM was not designed for this type of sample. We also omitted the uncorrected datasets based on SHERM’s instructions to correct for bias fields prior to running SHERM. In our preliminary tests, the maximally stable extremal region (MSER) selections were not able to survive the default convexity threshold of 0.85 on our test datasets. We lowered the threshold to 0.7 to produce better results, which was an approach also taken by Hsu et al. (2020).

##### 3D-PCNN

We applied 3D-PCNN (Chou et al., 2011) to the N4-corrected *in vivo* isotropic, *in vivo* anisotropic, and *ex vivo* datasets. Because the voxel resolutions of our images were not magnified, we set 3D-PCNN’s zoom factor parameter to 1 and modified the radii of its structural element to 5 for lower-resolution images (*≥* 100 µm in the dimension of the highest resolution) and 7 for higher-resolution images (*<* 100 µm in the dimension of the highest resolution), as per 3D-PCNN’s instructions.

##### RATS

We applied RATS to the N4-corrected test datasets only, as per the pre-processing methods described by the RATS developers (Oguz et al., 2014). We converted the resulting surface files, which RATS outputs in VTK polydata (VTP) format, into volumetric binary masks using functions available in 3DSlicer (Kikinis et al., 2013) as suggested by the RATS usage instructions.

##### rBET

We evaluated rBET (Wood et al., 2013) using all six test datasets. We specified an additional argument in rBET to set the brain radius to 5 mm to match mouse brain dimensions.

##### antsBrainExtraction

AntsBrainExtraction is an atlas-based method that requires a brain MRI template with an intact skull and a corresponding probability map of the brain region. We generated six separate MRI templates and sets of corresponding brain probability maps using the images and masks from the training datasets (see App. B for more details). We only evaluated antsBrainExtraction on the uncorrected images because the antsBrainExtraction workflow includes its own N4 bias field correction. We note that prior to running these steps, we modified the image header information to increase the voxel dimension sizes by a factor of 10 to approximate human MRI resolutions.

#### 2.9.2 Statistical Comparison

We evaluated the segmentation results for each of the methods using Dice and HD95 measures computed relative to the manually-edited data. We then performed two-tailed Welch’s t-tests to determine whether the results were statistically different from those of Mouse Brain Extractor. As described above, some methods were not run on all datasets because of the limitations of those methods. Additionally, 3D-PCNN failed to complete processing for four images, and DeepBrainIPP produced empty masks that did not identify any brain matter for 11 images. We ommitted these methods that produced an incomplete set of masks for a particular dataset from the subsequent analysis.

## 3 Results

We performed the processing steps as described in Section 2. The models for Mouse Brain Extractor, nnU-Net, and SwinUNETR were trained on multiple computers, each of which had either an NVIDIA Titan RTX or an NVidia GeForce RTX 4090 graphical processing unit (GPU) that was used for training. Each GPU had 24GB memory. We applied the trained Mouse Brain Extractor, SwinUNETR, and nnU-Net models and the other seven algorithms (antsBrainExtraction, RATS, rBET, SHERM, 3D-PCNN, DeepBrainIPP, RodentMRISkullStripping) to the test datasets detailed in Section 2.5.3 as summarized in Table 3. All methods completed without processing errors, with the exception of 3D-PCNN, which failed to finish on four of the *in vivo* anisotropic MRI. DeepBrainIPP failed to detect any brain matter in six uncorrected and five N4-corrected MRIs from the *in vivo* anisotropic test dataset.

We computed Dice coefficients and HD95 measures between each automated segmentation result and the corresponding manually-edited mask. The distributions of the Dice and HD95 measures for all methods are shown in Figs. 3 and 4, respectively. 3D-PCNN and DeepBrainIPP are omitted from the plots for the *in vivo* anisotropic test data results in Figs. 3D and 4D because neither method generated a complete set of brain segmentations for those datasets.

**Figure 3:**
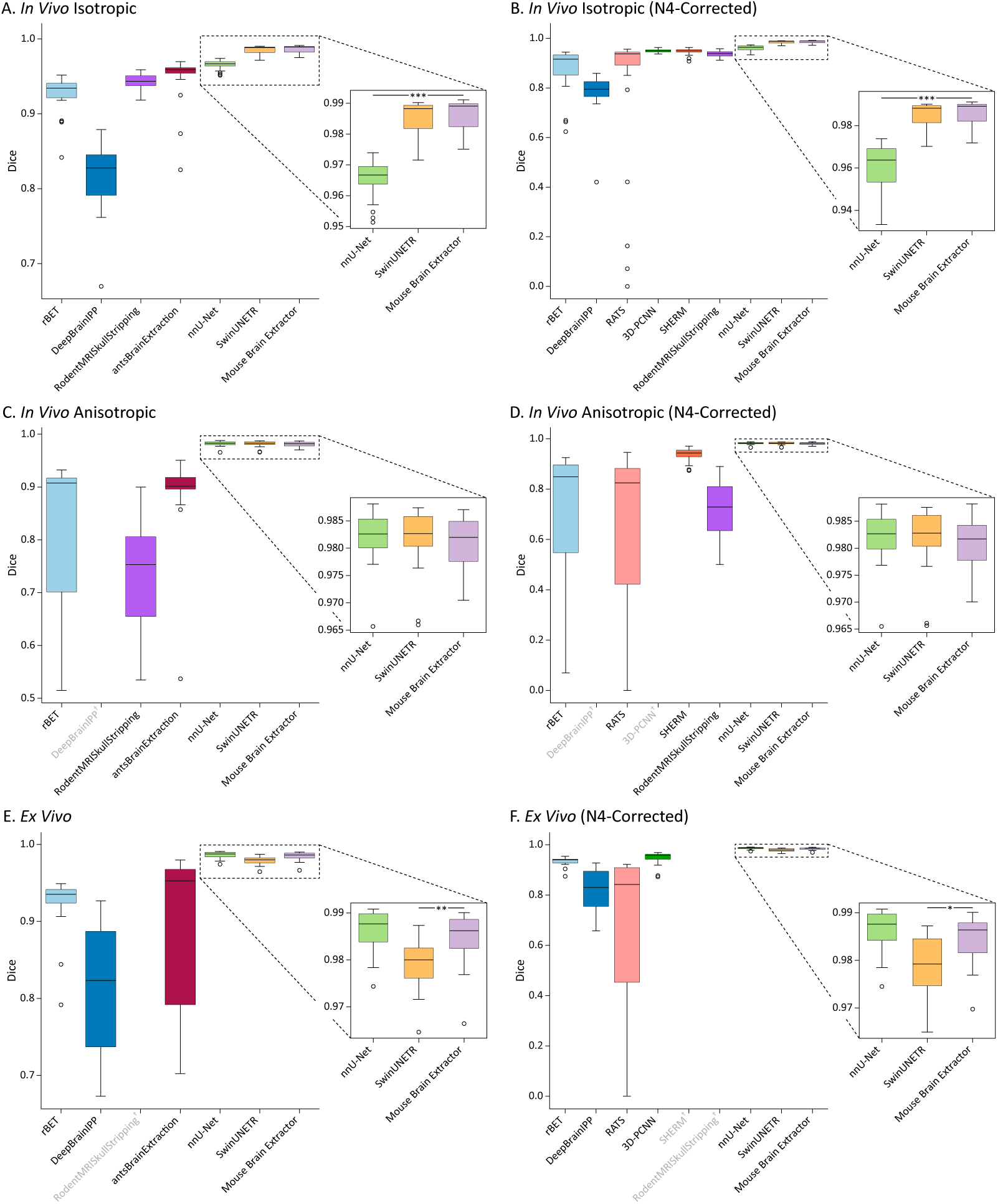
Dice Results. Box plots of the Dice similarity coefficients for the methods compared, applied to: A) uncorrected *in vivo* isotropic data; B) N4-corrected *in vivo* isotropic data; C) uncorrected *in vivo* anisotropic data; D) N4-corrected *in vivo* anisotropic data; E) uncorrected *ex vivo* data; F) N4-corrected *ex vivo* data. Higher Dice values indicate a better segmentation overlap and range between 0 and 1. We performed pairwise statistical comparisons on the Dice scores of our method Mouse Brain Extractor against those of other methods using two-tailed Welch’s t-tests. NnU-Net and SwinUNETR were the most competitive against Mouse Brain Extractor. We thus display for each plot an expanded view of nnUNet, SwinUNETR, and Mouse Brain Extractor. The significance bars and asterisks are shown only for the subplot (*p *<* 0.02; **p *<* 0.01; ***p *<* 5 × 10^−11^). Mouse Brain Extractor consistently outperformed the rest of the methods (rBET, DeepBrainIPP, RodentMRISkullStripping, antsBrainExtraction, RATS, 3D-PCNN, SHERM) across all datasets. †Greyed text indicates methods were not evaluated for a particular dataset (see Sec. 2.8 and App. B for details).

**Figure 4:**
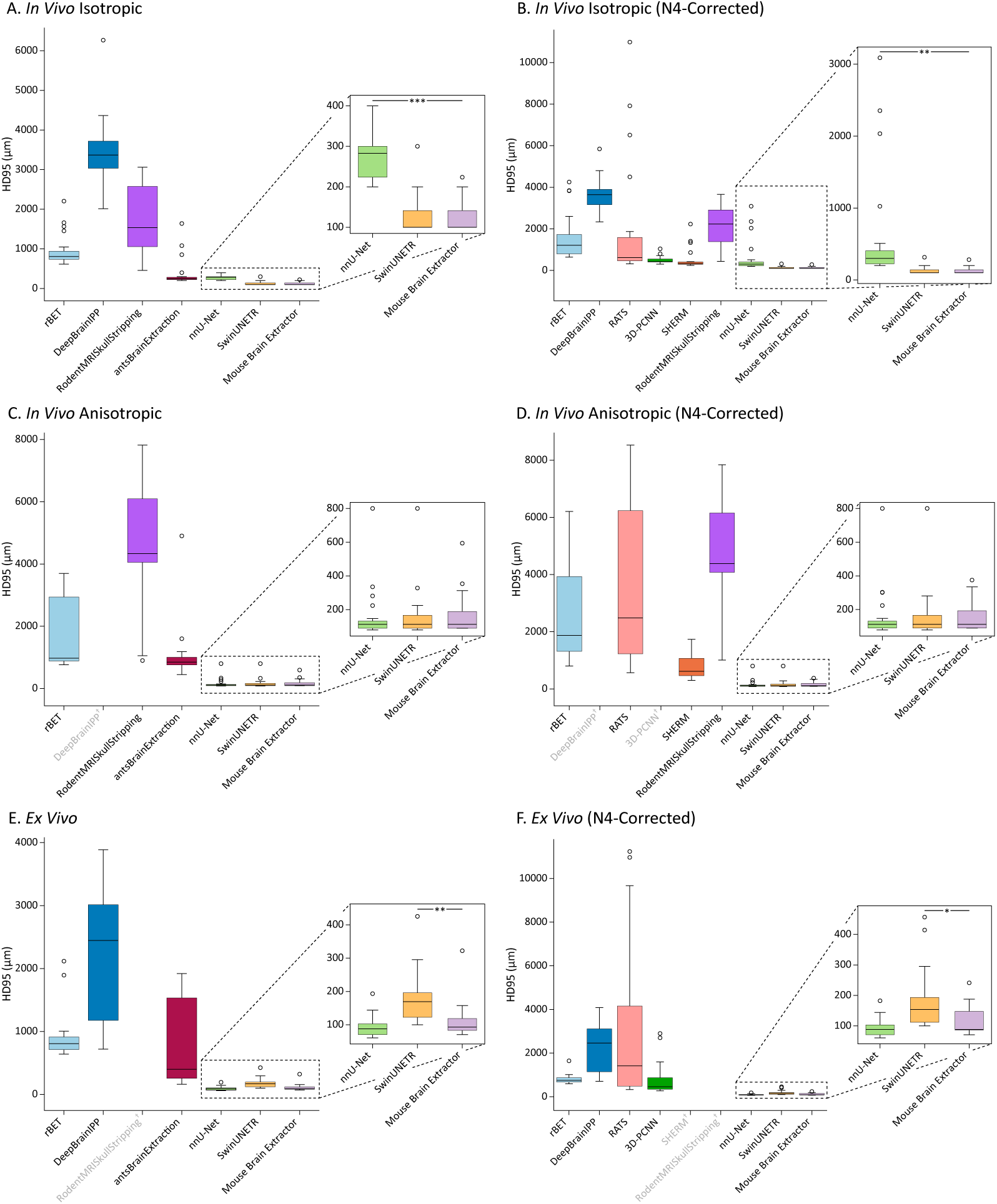
HD95 results. Box plots of the 95th percentile Hausdorff distance results for the methods compared, applied to: A) uncorrected *in vivo* isotropic data; B) N4-corrected *in vivo* isotropic data; C) uncorrected *in vivo* anisotropic data; D) N4-corrected *in vivo* anisotropic data; E) uncorrected *ex vivo* data; F) N4-corrected *ex vivo* data. Smaller distance measures indicate a closer proximity of segmentation boundaries and are measured in µm. We conducted pairwise two-tailed Welch’s t-tests on the HD95 measures of Mouse Brain Extractor and those of every other method. NnU-Net and SwinUNETR were the most competitive against Mouse Brain Extractor, and their results are shown in expanded views. We show significance bars and asterisks only for these subplots (*p *<* 0.02; **p *<* 0.01; ***p *<* 5 × 10^−11^). Mouse Brain Extractor achieved statistically lower HD95 measures on the rest of the methods (rBET, DeepBrainIPP, RodentMRISkullStripping, antsBrainExtraction, RATS, 3D-PCNN, SHERM) on all datasets. †Greyed text indicates methods were not evaluated for a particular dataset (see Sec. 2.8 and App. B).

We performed two-tailed Welch’s t-tests (Welch, 1947) on the Dice scores and the HD95 measures of Mouse Brain Extractor against those of the other methods. As shown in the box plots in Figs. 3 and 4, nnU-Net and SwinUNETR, and Mouse Brain Extractor had results that were competitive with each other, while the results for the remaining algorithms were worse in general. Mouse Brain Extractor’s average Dice and HD95 measures were statistically better than those of antBrainExtraction, RATS, rBET, SHERM, 3D-PCNN, DeepBrainIPP, and RodentMRISkullStripping for all test datasets (Dice: 10^−25^ *<* p *<* 10^−3^; HD95: 10^−17^ *<* p *<* 10^−3^). For the *in vivo* isotropic datasets, antsBrainExtraction, 3D-PCNN, SHERM and RodentMRISkullStripping also performed competitively against nnU-Net. We note that for RATS and RodentMRISkullStripping, we also performed comparisons excluding the coronal slices past the cerebellum, because these methods include parts of the spinal cord in the brain mask by design. Doing so provided small improvements in the Dice and HD95 measures. However, this did not change their overall competitiveness, and their evaluation metrics were still worse statistically than those of Mouse Brain Extractor.

We therefore focus primarily on the results of nnU-Net, SwinUNETR, and Mouse Brain Extractor. The Dice and HD95 results for these three methods are shown in expanded views for each plot in Figs. 3 and 4 to facilitate interpretation. Mouse Brain Extractor produced average Dice coefficients and HD95 measures that were better, with statistical significance, than those of nnU-Net for both sets of *in vivo* isotropic data (Figs. 3AB and 4AB). For the uncorrected *in vivo* isotropic data, Mouse Brain Extractor achieved an average Dice coefficient of 0.9863±0.0048 and an average HD95 of 120.5±33.5 µm. These results were significantly better than nnU-Net’s mean Dice coefficient of 0.9654±0.0061 (p = 7.55 × 10^−16^) and mean HD95 measure of 266.0±52.0 µm (p = 2.44 × 10^−13^). Similarly, for the N4-corrected *in vivo* isotropic data, Mouse Brain Extractor yielded higher Dice scores (0.9863±0.0051; p = 1.61 × 10^−11^) and lower HD95 metrics (124.9±42.3 µm; p = 7.5 × 10^−3^) than nnU-Net (Dice: 0.9606±0.0106; HD95: 606.1±766.4 µm). Mouse Brain Extractor performed comparably to SwinUNETR and did not differ statistically on either the uncorrected *in vivo* isotropic data (Dice: 0.9855±0.0052, p = 0.5807; HD95: 127.0±46.1 µm, p = 0.5954) or the N4-corrected *in vivo* isotropic data (Dice: 0.9854±0.0052, p = 0.5727; HD95: 129.5±48.5 µm, p = 0.7371).

For the uncorrected and N4-corrected *in vivo* anisotropic datasets (Fig. 3C&D), the average Dice scores for nnU-Net and SwinUNETR were slightly higher than those of Mouse Brain Extractor, but these differences did not reach statistical significance. The average HD95 measures (Fig. 4C&D) for nnUNet were lower than those of Mouse Brain Extractor, but again these results were not statistically significant. Specifically, for the uncorrected *in vivo* anisotropic data, Mouse Brain Extractor achieved an average Dice coefficient of 0.9808±0.0048 and an average HD95 of 165.0±112.7 µm, which were not significantly different from the corresponding results of nnU-Net (Dice: 0.9821±0.0045, p = 0.2906; HD95: 157.1±138.9 µm, p = 0.8199) or SwinUNETR (Dice: 0.9816±0.0052, p = 0.5277; HD95: 179.7±181.6 µm, p = 0.7222). For the N4-corrected *in vivo* anisotropic data, Mouse Brain Extractor achieved an average Dice coefficient of 0.9806±0.0047 and average HD95 of 162.8±92.3 µm, which again did not differ significantly from nnU-Net (Dice: 0.9822±0.0045, p = 0.2137; HD95: 156.2±138.4 µm, p = 0.8392) or SwinUNETR (Dice: 0.9817±0.0054, p = 0.4191; HD95: 175.7±181.5 µm, p = 0.7432).

For the *ex vivo* datasets, Mouse Brain Extractor produced average Dice coefficients and average HD95 measures that were better statistically than those of SwinUNETR (Figs. 3EF and 4EF). For the uncorrected *ex vivo* data, Mouse Brain Extractor produced a mean Dice score of 0.9844±0.0059 and a mean HD95 measure of 111.0±55.4 µm, which were significantly better than SwinUNETR’s mean Dice score (0.9791±0.0056; p = 7.65 × 10^−3^) and mean HD95 measure (179.9±76.7 µm; p = 3.18 × 10^−3^). Similarly, for the N4-corrected *ex vivo* data, Mouse Brain Extractor yielded higher mean Dice (0.9842±0.0054; p = 0.0112) and HD95 measures (114.3±44.3 µm; p = 0.0101) than SwinUNETR (Dice: 0.9793±0.0059; HD95: 182.8±98.4 µm). Mouse Brain Extractor performed comparably to nnU-Net and did not differ significantly on either the uncorrected data (Dice: p = 0.3260; HD95: p = 0.3182) or the N4-corrected data (Dice: p = 0.2603; HD95: p = 0.1469).

We additionally assessed whether N4-correction significantly altered the automated methods’ performances. We computed two-tailed Welch’s t-tests on the evaluation metrics derived from the predicted segmentations generated using uncorrected and N4-corrected data. Interestingly, N4-correction negatively impacted nnU-Net’s performance on the *in vivo* isotropic data, for which nnU-Net rendered a significantly worse HD95 measure in the N4-corrected data compared to the uncorrected data (p = 0.0496). No other statistically significant difference was observed across the other methods and dataset types.

Although Dice scores and HD95 provide quantitative measures of segmentation similarities, they do not convey spatial information about the specific regions where the methods exhibited accurate or inaccurate performance. We thus show in Fig. 5 examples of individual segmentation outputs produced by nnUNet, SwinUNETR, and Mouse Brain Extractor, compared individually with the manually labeled masks. The subjects shown correspond to the images that resulted in the best and worst Dice scores for Mouse Brain Extractor on the uncorrected data; for consistency, we show the same subjects and slices for all three methods. As observed in the figure, the boundaries in the superior frontal regions were labeled correctly in general by all three methods across all datasets. In contrast, each of these methods often labeled the ventral regions of the brain incorrectly, possibly because of attenuated signals caused by severe inhomogeneity fields. Each method commonly mis-segmented the paraflocculi in the *in vivo* anisotropic dataset, perhaps because of substantial partial volume effects caused by large slice thicknesses. In the *ex vivo* data, only Mouse Brain Extractor incorrectly labeled the non-brain areas near the cerebellum, specifically, neck muscles. This may be due to the unclear boundary of the brain in this area. We note that applying N4 correction improved Mouse Brain Extractor’s segmentation of this area.

**Figure 5:**
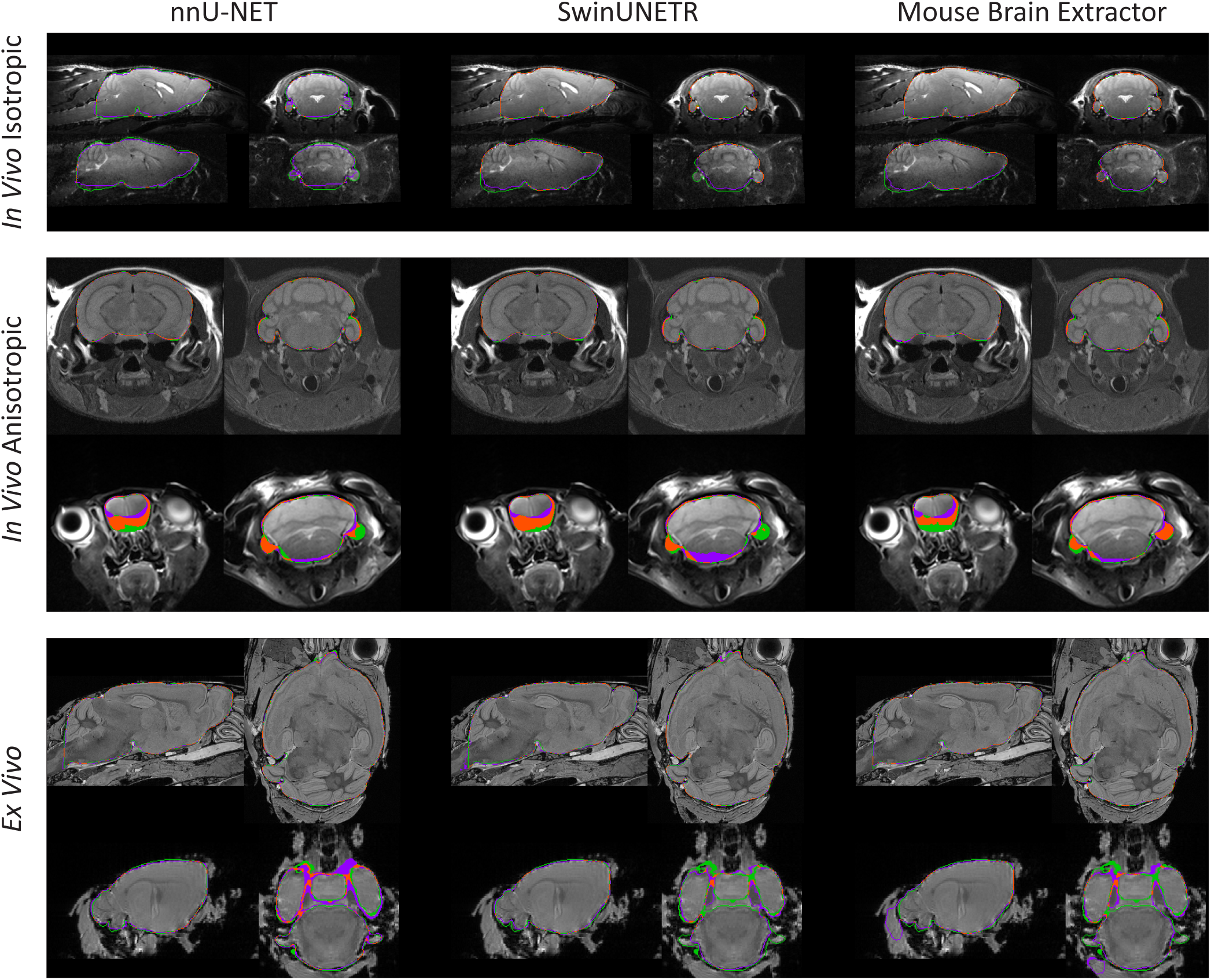
Examples of individual brain segmentations generated by nnU-Net, SwinUNETR, and Mouse Brain Extractor. Green pixels indicate the boundary voxels identified only by the manually-edited brain masks, while purple pixels indicate boundary voxels labeled only by the automated methods. Orange pixels indicate agreement between the manual and automated segmentation methods. For each dataset, the best and worst cases (top and bottom rows, respectively) were selected based on Mouse Brain Extractor’s Dice scores. All segmentation results and MRI slices shown are from the uncorrected test datasets. All three methods perform well near the superior frontal portion of the brain but had reduced accuracy in the ventral regions as well as in the paraflocculus. We note that when the boundaries are parallel to the plane of section, the contour lines appear thicker. Shown are: *in vivo* isotropic images from (top) a healthy control male from the EAE cohort and (bottom) a healthy male from the NeAT cohort; *in vivo* anisotropic images from (top) a B6-CC61 heterozygous male from the *Chd8* cohort and (bottom) a sham surgery female from the TBI cohort; and *ex vivo* images from (top) an EAE-induced female from the EAE cohort and (bottom) a healthy female from the EAE cohort.

We also created error maps to visualize where the methods most commonly fail. Following the approach we previously applied when evaluating skull-stripping methods in human MRI (Shattuck et al., 2009), we created spatial maps of the false positive and false negative errors relative to the manual delineations. We first applied ANTs nonlinear registration (Avants et al., 2011) to each image in each dataset to produce a spatial transform to the Mortimer Space Atlas (MSA50; Meyer et al., 2017). For each method being evaluated, we computed image arrays of false positive and false negative values by comparing its output masks with the manually-edited masks at each voxel. We resampled these false positive and false negative images to the MSA50 space using the ANTs transforms and linear interpolation, and then averaged the maps across each dataset to produce separate mean false positive and mean false negative images. These volumes thus represent the frequency of error at each voxel in the atlas space. We then summed the average error maps along the x-axis to produce sagittal views of the error densities.

Figures 6 and 7 show the spatial maps of false positives and false negatives, respectively, for nnU-Net, SwinUNETR, and Mouse Brain Extractor computed for the uncorrected versions of the *in vivo* isotropic, *in vivo* anisotropic, and *ex vivo* data. Results from the N4-corrected data produced similar maps and are omitted for brevity. As seen in Fig. 6, all three methods showed relatively high false positives in the ventral region where the trigeminal nerves reside, as well as in the posterior (brainstem-spinal cord) boundary, indicating that the methods often mislabeled these structures as brain matter. In particular, SwinUNETR’s errors in the *ex vivo* data were mainly in the brainstem-spinal cord boundary, where SwinUNETR mislabeled the spinal cord as brain. NnU-Net also showed higher false positives in the regions posterior to the cerebellum in the isotropic data, but nnU-Net’s errors were distributed more spatially across the whole brain.

**Figure 6:**
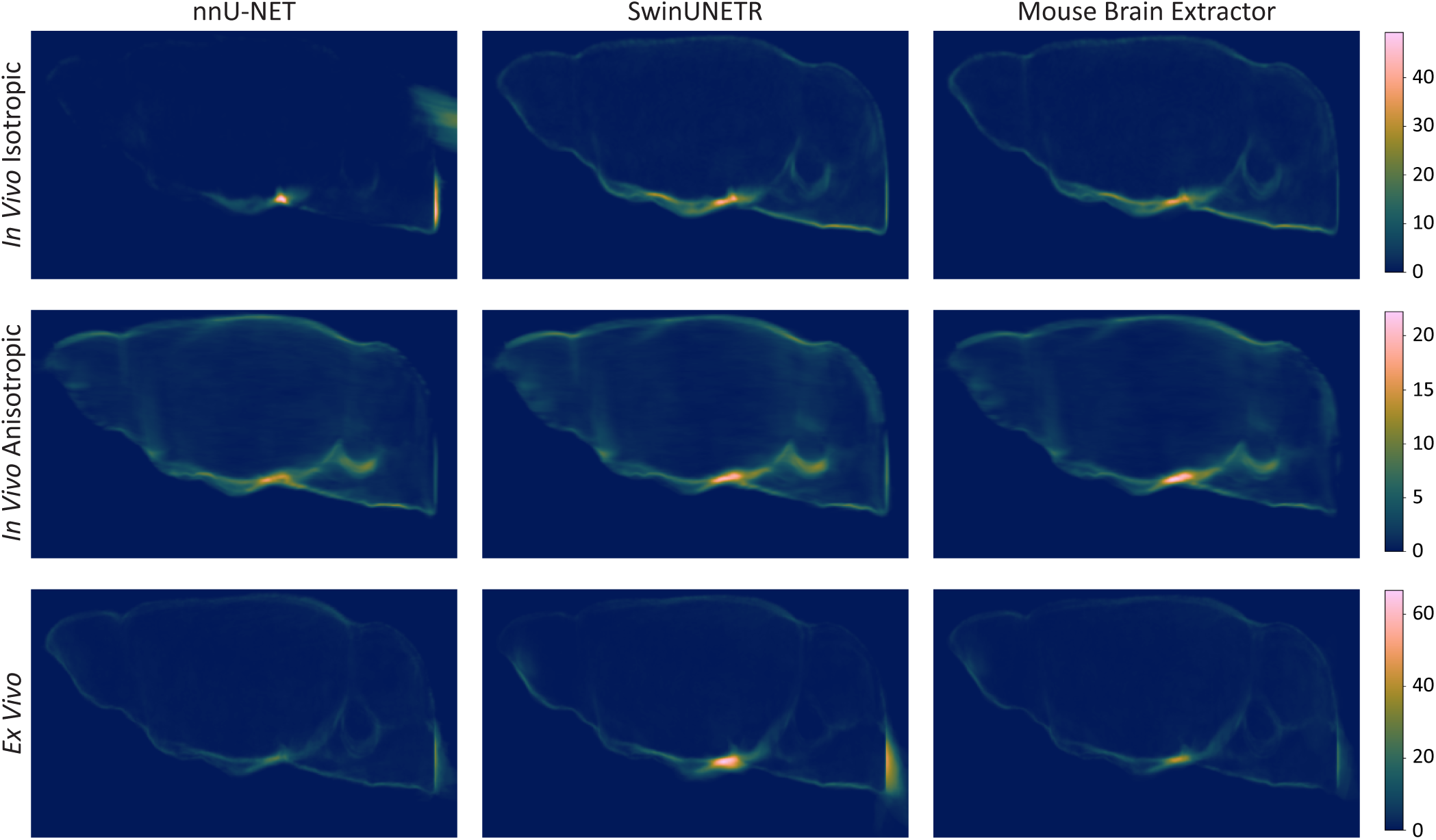
Average counts of false positive voxels for nnU-Net, SwinUNETR, and Mouse Brain Extractor. The images show where non-brain voxels were commonly mislabeled as brain. Each image displays the average false positive count along the x-axis. For each method and dataset, the false positive maps were registered non-linearly to the MSA50 atlas (Meyer et al., 2017), resampled to the atlas space, summed along the x-axis, and averaged over subjects. In general, higher average false positive counts were concentrated in the trigeminal nerve and spinal cord regions compared to other areas.

**Figure 7:**
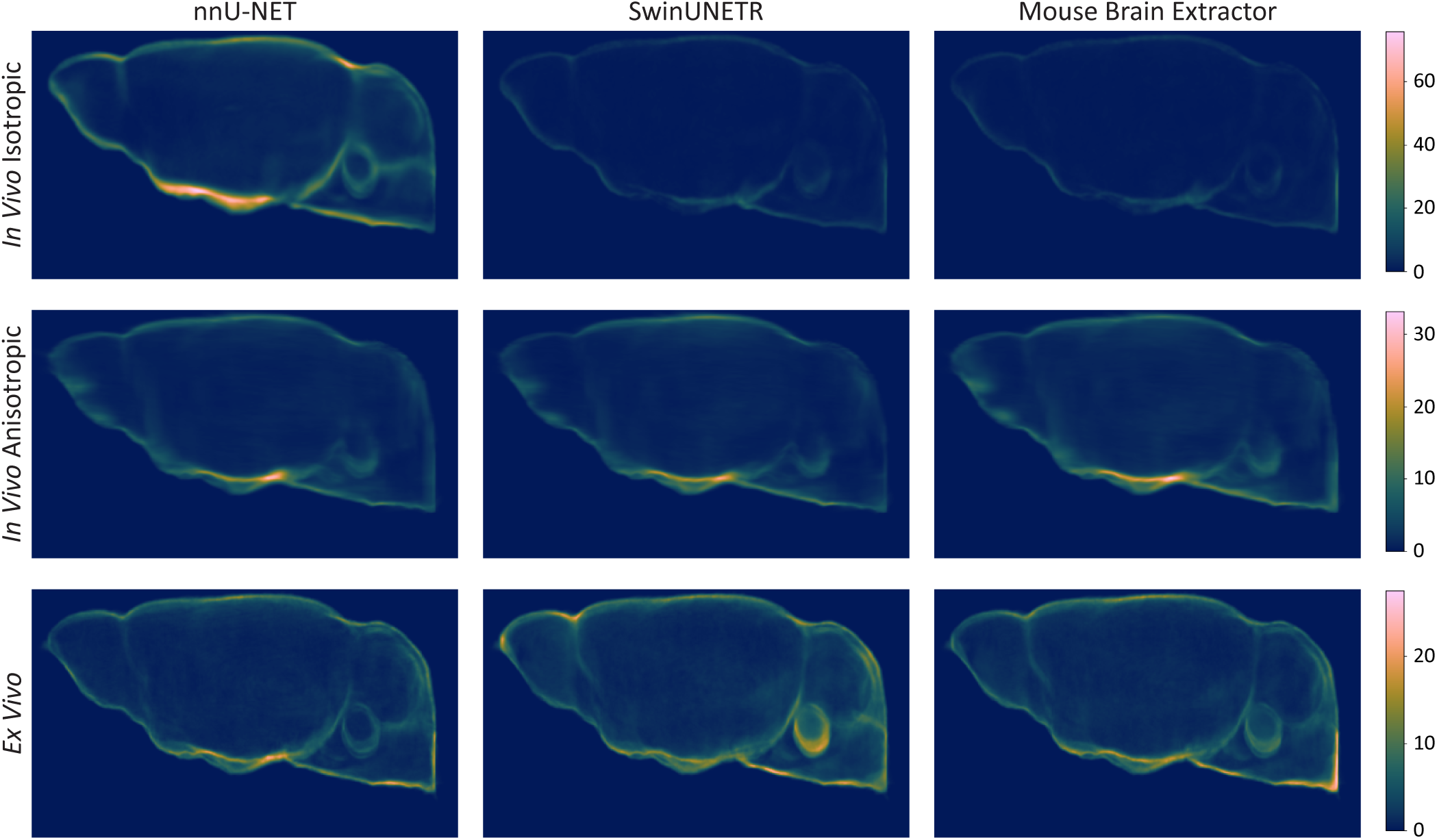
Average counts of false negative voxels for nnU-Net, SwinUNETR, and Mouse Brain Extractor. The images show areas where brain voxels were commonly mislabeled as non-brain. For each method and dataset, the false negative maps were aligned to the MSA50 atlas (Meyer et al., 2017) using nonlinear registration, resampled to the template space, summed along the x-axis, and averaged over subjects. Common regions of false negatives included the trigeminal nerve, the spinal cord, and the paraflocculus.

In the average false negative maps (Fig. 7), more errors can be seen in the regions near the trigeminal nerves in all datasets for all three methods. NnU-Net’s *in vivo* isotropic segmentation outputs produced a higher false negative count around the brain boundaries, indicating that there was frequent misidentification of brain matter in these areas. The ventral regions in particular showed higher false negatives. For the *ex vivo* data, SwinUNETR occasionally mislabeled the parafloccular nodules as non-brain matter, whereas Mouse Brain Extractor mislabeled parts of the spinal cord as non-brain.

## 4 Discussion

We have presented Mouse Brain Extractor, a method for segmenting the brain from whole-head mouse MRI data. Mouse Brain Extractor builds upon SwinUNETR (Hatamizadeh et al., 2021) by incorporating a new variation of absolute positional encoding called Global Positional Encoding. Our position embeddings are based on a shared coordinate frame that encompasses the whole input image. As a result, Mouse Brain Extractor performed competitively without needing to resample the input images or the segmentation outputs. Furthermore, to increase our model’s generalizability, we curated a heterogeneous collection of previously-acquired mouse MRI data, encompassing a variety of populations, sexes, ages, animal preparations, genotypes, and strains. These datasets also featured diverse MRI contrasts acquired using an array of scanner types and protocols. We generated corresponding brain masks by manually editing and minimally postprocessing an initial set of brain segmentations, which we used as ground truth labels. Based on our evaluations, Mouse Brain Extractor achieved Dice coefficients of approximately 0.98 and HD95 measures of approximately 100 to 200 µm when assessed against the ground truth labels. Our proposed method consistently outperformed rBET, DeepBrainIPP, RodentMRISkullStripping, antsBrainExtraction, RATS, 3D-PCNN, and SHERM across all datasets with statistical significance.

In comparison with nnU-Net and SwinUNETR, two state-of-the-art deep learning methods, Mouse Brain Extractor also performed competitively. Mouse Brain Extractor produced significantly better Dice and HD95 values than nnU-Net on *in vivo* isotropic resolution datasets and than SwinUNETR on *ex vivo* datasets. When considering the mean values, Mouse Brain Extractor achieved a Dice score that was approximately 0.02 higher than that of nnU-Net in the *in vivo* isotropic dataset. In the *in vivo* anisotropic resolution datasets, Mouse Brain Extractor performed comparably to nnU-Net and SwinUNETR and did not differ statistically in either Dice or HD95 measures. For the *ex vivo* data, Mouse Brain Extractor resulted in Dice scores that were approximately 0.005 higher than those of SwinUNETR. We note that although the Dice similarity improvements appear modest, the brain segmentations were composed of a large number of voxels and a relatively smooth object of interest. This means that the the scores for the best methods were highly compressed near the maximum value of the Dice similarity coefficient. As the segmentation sizes become larger, the incremental increases in the segmentations produce smaller changes in the resulting Dice scores. For example, consider two cubic sets of segmented voxels *T* and *U*, where *T* is a cube of size (*N* + 2) *×* (*N* + 2) *×* (*N* + 2) and *U* is a cube of size *N × N × N* comprising the interior voxels of *T*. In this case, the Dice coefficient can be simplified to 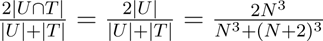. If *N* = 10, the Dice score is approximately 0.733. However, if the sizes of the cubes are increased such that *N* = 100, the Dice score becomes approximately 0.970. For larger objects, single-voxel boundary shifts have a reduced effect on the Dice score. Brain segmentation methods that perform well often differ primarily near the boundary voxels, but match in the interior of the brain. Improvements in detecting the boundary accurately can thus appear less impactful. For cases with larger segments, boundary measures can provide more informative quantitative metrics than Dice scores. This can be seen in the HD95 measures, which were on the order of 110 µm for Mouse Brain Extractor and 180 µm for SwinUNETR. Here, the discrepancies in the distances are more noticeable compared to the differences in the Dice scores.

We observed similar statistical significance in the evaluation of segmentation results generated from the N4-corrected images by nnU-Net, SwinUNETR, and Mouse Brain Extractor. We also note that all three methods were robust to bias field artifacts and N4-correction, with the exception of one case from nnU-Net. In the *in vivo* isotropic data, nnU-Net’s HD95 increased by 340 µm in the N4-corrected data compared to the uncorrected data. One possible reason for the worse performance is that N4-correction could have inaccurately transformed the image intensities. The bias field artifacts were particularly severe in the *in vivo* isotropic dataset, thus N4-correction could have increased the effect of the artifact by pushing the intensity image values in the wrong direction. This would have caused the existing hyperintense voxels to become brighter and hypointense areas to become darker. Mouse Brain Extractor was less sensitive to the effects of N4-correction and yielded HD95 measures that were on the order of 120 µm for both the uncorrected and N4-corrected data. Additionally, N4-correction may not be a necessary pre-processing step for Mouse Brain Extractor or for SwinUNETR because they performed comparably on both uncorrected and N4-corrected data, with no statistical significance.

In general, there were some anatomical areas that nnU-Net, SwinUNETR, and Mouse Brain Extractor commonly mislabeled. One of these was the ventral region of the brain. There were substantial bias field artifacts in the *in vivo* datasets, which produced signals that were often attenuated in the ventral regions. This may have made it more challenging for the automated methods to detect edges. In particular, nnUNet often mis-segmented these areas in the *in vivo* isotropic datasets. NnU-Net’s accuracy may have suffered in this case due to the lack of global spatial context, which was limited because of its 2D input or convolutional architecture. Conversely, Mouse Brain Extractor, which incorporated 3D inputs and Transformer blocks, produced far fewer errors in these same regions. Field inhomogeneity can also make it difficult to identify the paraflocculus. This was especially challenging in the *in vivo* anisotropic datasets, because the inter-plane slice thicknesses were much larger compared to the other datasets. The paraflocculi are small in size, thus the partial volume effect greatly exacerbated the blurring of their boundaries.

Other common areas where segmentation errors occurred included the trigeminal nerve boundary and the brainstem-spinal cord boundary. These boundaries do not exhibit distinct physical features and require contextual information from the surrounding anatomy to be identified correctly. According to the convention of our manual delineation protocol, the trigeminal nerve is included in the brain mask when the peduncles have appeared in the coronal view. Similarly, our protocol excludes the spinal cord from the brain mask when the cerebellum is not present in the coronal view. The automated methods may have difficulty delineating these boundaries because the algorithms based their boundaries on local spatial patterns and contrasts, whereas the human rater followed a protocol that determines a delineation, in part, by the appearance of anatomical structures in a slice that is dependent on a specific scanning orientation (e.g., the coronal plane). Such errors were observed frequently in SwinUNETR’s segmentation results for the *ex vivo* datasets, which included images with large sizes and multiple image resolutions. However, with the inclusion of Global Positional Encoding in our Mouse Brain Extractor method, performance in these regions improved. This suggests that the GPEs may have provided the model with additional spatial context regarding the subimage’s location relative to the whole image. In contrast, in the case of the *in vivo* anisotropic data, which consisted of larger subimage sizes and smaller input image volumes, SwinUNETR performed competitively, possibly because more of the image could be captured in the subimage.

A key strength of our approach is its ability to perform well across multiple different types of acquisitions and mouse brain models. In addition to the variety of image quality and resolutions, the datasets used for this study encompassed a wide range of brain disorders with distinct anatomical differences. For example, the TBI dataset contained mice with abnormal brain deformations that were markedly distinct from the macrocephalic *Chd8* mice. Despite these differences in anatomy and image appearance, Mouse Brain Extractor was consistently able to segment the brains accurately.

One limitation of our study is the lack of T1-weighted contrasts in the *in vivo* MRI datasets, which makes our method less generalizable to these types of data. Another limitation is the longer inference time compared to nnU-Net. This is primarily because of an increased number of multiply-add operations and model parameters in Mouse Brain Extractor and the calculation of the global positional encodings. This makes our method slower than nnU-Net and SwinUNETR by approximately 3-fold and 1.2-fold, respectively. NnU-Net contains on the order of 30 million parameters and requires approximately 200 billion floating point operations (FLOPs) to infer a single brain mask for an input MRI. SwinUNETR and Mouse Brain Extractor each have on the order of 40 billion parameters and require approximately 700 billion FLOPs for the neural network to infer a single brain mask. Additionally, the generation of the 48-element global positional encodings in the Mouse Brain Extractor, which are computed immediately prior to model inference, further increases the number of FLOPs and the computation time by approximately 20-fold compared to SwinUNETR.

We plan to extend the present work in multiple ways. We are currently incorporating the brain segmentation software into a larger, more comprehensive analysis framework for performing group studies on preclinical images. We also plan to train and test these tools on rat brain MRI to expand its application in preclinical rodent imaging. Finally, we are exploring ways to improve the computational efficiency of the methods.

## Ethics Statement

The MRI data used in this study were acquired previously in separate studies (Hubbard et al., 2021; Itoh et al., 2023; MacKenzie-Graham et al., 2009; Meyer et al., 2017), as part of ongoing studies of specific mouse models (Tabbaa et al., 2023), or retrieved from an open-access repository (Y. Ma et al., 2008). All studies from which we retrieved data were performed in accordance with the National Institutes of Health Guide for the Care and Use of Laboratory Animals and appropriate Institutional Animal Care and Use Committees (IACUC).

## Data and Code Availability

The Python source code for Mouse Brain Extractor is released under the the GNU General Public License v2.0 only (GPL-2.0-only) and is available from our GitHub repository: https://github.com/MouseSuite/ MouseBrainExtractor. We have also released Docker and Singularity/Apptainer images that contain the pre-trained weights and all software dependencies necessary to run Mouse Brain Extractor. Instructions for use are provided on our GitHub page.

The MRI data used from the MRM NeAt dataset are publicly available the NeAT GitHub repository: https://github.com/dama-lab/mouse-brain-atlas/tree/master/NeAt/in vivo. The remaining MRI data used in this paper are not publicly available. MRI data for the EAE, aging, and TBI cohorts are available upon reasonable request, which may be sent to the corresponding author. Original MRI data for the *Chd8* dataset will be available upon reasonable request once the neuroanatomical strain differences are reported in a separate publication. Requests for access will be considered by the relevant coauthors in accordance with necessary data sharing agreements or other institutional requirements.

## Author Contributions

**Yeun Kim:** Conceptualization, Data curation, Software, Formal analysis, Investigation, Methodology, Resources, Visualization, Writing - original draft, Writing - review & editing; **Haley Hrcnir:** Data curation, Resources, Writing - review & editing; **Cassandra Meyer:** Data curation, Resources, Writing - review & editing; **Manal Tabbaa:** Resources, Writing - review & editing; **Rex Moats:** Data curation, Resources, Writing - review & editing; **Pat Levitt:** Resources, Writing - review & editing; **Neil Harris:** Data curation, Resources, Writing - review & editing; **Allan MacKenzie-Graham:** Data curation, Funding acquisition, Methodology, Resources, Writing - original draft, Writing - review & editing; **David Shattuck:** Conceptualization, Software, Funding acquisition, Methodology, Project administration, Resources, Supervision, Visualization, Writing - original draft, Writing - review & editing.

## A MRI data summary

In this section, we describe the imaging acquisition procedures that were used to acquire the MRI data used in our study (see Table 4) and the demographic details for the mice that were scanned. A subset of these data (see Table 5) were used to evaluate the different methods compared in this paper. All imaging data were collected previously as parts of other studies (Hubbard et al., 2021; Itoh et al., 2023; MacKenzie-Graham et al., 2009; Meyer et al., 2017), were acquired as part of ongoing studies of specific mouse models (Tabbaa et al., 2023), or were retrieved from a publicly available open-access repository (Y. Ma et al., 2008).

**Table 4:** MRI description for the data used in this study. In total, our study gathered datasets that were scanned using eight distinct pulse sequences at five different MR scanners. All datasets except for the *in vivo* isotropic resolution images had varying degrees of resolutions. Voxel aspect ratio is the ratio of the maximum to the minimum voxel edge length. X-, y-, and z-axis refer to a right-anterior-superior orientation, where x-axis represents left/right, y-axis represents rostral/caudal, and z-axis represents dorsal/ventral.

| Dataset | Cohort | Voxel Aspect Ratio [min,max] | Scanner | Sequence | Image resolutions ($\mu\text{m} \pm \text{sd}$) | | | Image dimensions [min, max] | | |
| --- | --- | --- | --- | --- | --- | --- | --- | --- | --- | --- |
|  |  |  |  |  | x-axis | y-axis | z-axis | x-axis | y-axis | z-axis |
| *in vivo* isotropic | EAE<br>Meyer et al., 2017 | [1, 1] | 7 T Bruker;<br>11.7 T Biospec<br>Bruker | T2 RARE (TR=3500 ms;<br>TE<sub>eff</sub>=32 ms; ETL=16) | 100.0 ± 0.0 | 100.0 ± 0.0 | 100.0 ± 0.0 | [192, 192] | [256, 256] | [100, 100] |
| | Aging<br>Itoh et al., 2023 | [1, 1] | 7 T Bruker;<br>11.7 T Biospec<br>Bruker | T2 RARE (TR=3500 ms;<br>TE<sub>eff</sub>=32 ms; ETL=16) | 100.0 ± 0.0 | 100.0 ± 0.0 | 100.0 ± 0.0 | [192, 192] | [256, 256] | [100, 100] |
| | NeAt<br>Ma et al., 2008 | [1, 1] | 9.4 T Magnex/<br>Bruker | T2 spin echo (TR=400 ms;<br>TE=7.5 ms) | 100.0 ± 0.0 | 100.0 ± 0.0 | 100.0 ± 0.0 | [192, 192] | [256, 256] | [96, 96] |
| *in vivo* anisotropic | *Chd8*<br>Tabbaa et al., 2023 | [3.98, 6.70] | 11.7 T Biospec<br>Bruker | T2 turbo RARE | 56.1 ± 0.3 | 333.7 ± 68.4 | 56.0 ± 0.4 | [284, 320] | [48, 80] | [284, 320] |
| | TBI<br>Hubbard et al., 2021 | [8.57, 15.36] | 7 T Bruker/<br>Siemens<br>Clinscan | T2 turbo spin echo<br>(TR=2070 ms; TE=42 ms) | 80.4 ± 16.5 | 777.5 ± 13.1 | 80.4 ± 16.5 | [256, 384] | [18, 21] | [208, 312] |
| *ex vivo* | EAE<br>MacKenzie-Graham et al., 2009 | [1, 1.85] | 11.7 T Bruker | T2 RARE (TR=1500 ms;<br>TE<sub>eff</sub>=10 ms) | 46.2 ± 0.8 | 50 ± 0.0 | 37.5 ± 0.0 | [256, 512] | [512, 512] | [256, 256] |
| | | | 9.4 T Agilent | RF refocused spin echo | 43.0 ± 0.0 | 43.0 ± 0.0 | 43.0 ± 0.0 | [256, 256] | [512, 512] | [256, 256] |
| | | | 11.7 T Biospec<br>Bruker | T1 FLASH (TR=58.885 ms;<br>TE=7.713 ms); | 64.5 ± 0.0 | 64.5 ± 0.0 | 64.5 ± 0.0 | [154, 169] | [217, 272] | [127, 183] |
| | | | | T1 FLASH (TR=23.3 ms;<br>TE=4.38 ms) | 50 ± 0.0 | 50 ± 0.0 | 50 ± 0.0 | [244, 284] | [330, 389] | [159, 204] |

**Table 5:** Demographic information for the test datasets. A subset of images representative of each cohort was used to evaluate the methods. Voxel aspect ratio is the ratio of the maximum to the minimum voxel edge length and days-post-injury (DPI) indicates the number of days after brain injury or sham surgery.

| Dataset | Cohort | Strain | Genotype | Sex | Age at Scan (mos. ± sd) | Notes | Total |
| --- | --- | --- | --- | --- | --- | --- | --- |
| *in vivo* isotropic | EAE<br>Meyer et al., 2017 | 15 C57BL/6 | 15 WT | 11 F / 10 M | 4.91 ± 0.48 | 9 EAE / 12 healthy controls | 15 |
| | Aging<br>Itoh et al., 2023 | 6 C57BL/6 | 6 WT | 3 F / 3 M | 13.17 ± 7.94 | Aged 5 to 24 months | 6 |
| | NeAt<br>Ma et al., 2008 | 2 C57BL/6 | 2 WT | 0 F / 2 M | 3.25 ± 0.25 | | 2 |
| *in vivo* anisotropic | *Chd8*<br>Tabbaa et al., 2023 | 7 B6-CC61<br>5 B6-CC17 | 6 HET<br>6 WT | 6 F / 6 M | 10.68 ± 1.52 | | 12 |
| | TBI<br>Hubbard et al., 2021 | 16 C57BL/6 | 16 WT | 8 F / 8 M | 4.97 ± 2.21 | 8 injured / 8 sham<br>4 0-DPI / 4 30-DPI / 4 100-DPI / 4 166-DPI | 16 |
| *ex vivo* | EAE<br>MacKenzie-Graham et al., 2009 | 20 C57BL/6 | 20 WT | 17 F / 3 M | 4.66 ± 1.34 | 13 EAE / 7 healthy controls<br>4 SOD1<sup>-/-</sup><br>4 AAV-treated | 20 |

### A.1 *In vivo* experimental autoimmune encephalomyelitis (EAE) and aging data

The *in vivo* EAE and aging dataset we selected for our study consisted of 30 EAE-induced mice and 44 healthy controls. Twelve of the healthy control mice were scanned at the Small Animal Imaging Core (SAIC) at the Children’s Hospital Los Angeles (CHLA) using an 11.7 T BioSpec (Bruker Instruments, Billerica, MA) scanner. The remaining mice were scanned at the Ahmanson-Lovelace Brain Mapping Center (ALBMC) at UCLA using a 7 T Bruker (Bruker Instruments, Billerica, MA) scanner. Both cohorts were imaged using a rapid-acquisition with relaxation enhancement (RARE) sequence (TR/TE_eff_ = 3500/32 ms, ETL = 16, matrix = 256 *×* 192 *×* 100, voxel dimensions = 100 µm isotropic resolution) (Itoh et al., 2023; Meyer et al., 2017).

### A.2 MR microscopy (MRM) NeAt data

For our study, we used five mouse MRI (5M; age at scan 3.25 ± 0.25 months) from the publicly available MRM NeAt dataset (https://github.com/dama-lab/mouse-brain-atlas/tree/master/NeAt/in vivo), which includes 11 male T2w images, with ages ranging from 3 to 3.5 months old at the time of the scan (Y. Ma et al., 2008). We excluded six of these images from our study because of their poor quality. All MRM NeAt mouse MRI data were acquired using a 9.4 T Magnex horizontal bore magnet with an ADVANCE Bruker console (Bruker, Billerica, MA) with the following acquisition parameters: 145° flip angle; TE = 7.5 ms; TR = 400 ms; 100 *µ*m isotropic resolution (Ali et al., 2005; Bogdan & Joseph, 1990; DiIorio et al., 1995; Elster & Provost, 1993; J. Ma et al., 1996). Additional details regarding the MRM NeAt data can be found in (Y. Ma et al., 2008).

### A.3 *Chd8* data

All 36 mice from the *Chd8* cohort were acquired at SAIC using an 11.7 T Bruker BioSpin (Bruker Instruments, Billerica, MA) using two similar T2 sequences. Twenty-six mice (13F/13M; 13 HET/13WT; 12 B6-CC17/14 B6-CC61; age at scan 11.16 ± 1.1 months) were scanned using a T2 turbo RARE sequence (TE = 20 ms; TR = 2800 ms; 48 slices; field of view = 18 mm *×* 18 mm; in-plane matrix = 320 *×* 320, slice thickness 0.056 mm). Images from another 10 mice (5F/5M; 5 HET/5WT; 3 B6-CC17/7 B6-CC61; age at scan 2.03 ± 0.2 months) were acquired on the same scanner using a slightly different T2 turbo RARE sequence (TE = 20 ms; TR = 2250; 40 slices; field of view = 16 mm *×* 16 mm; in-plane matrix = 284 *×* 284, slice thickness 0.214 mm).

### A.4 Traumatic brain injury (TBI) data

The mice from the TBI study were scanned using a 7T Bruker/Siemens Clinscan scanner using a turbo spin echo sequence (TR = 2070 ms; TE = 42 ms; field of view = 40 mm *×* 0.7 mm *×* 40 mm; in-plane matrix = 384 *×* 512). Twelve consecutive slices were acquired near the site of controlled cortical impact (CCI) injury or sham surgery. More information on the imaging is available in (Hubbard et al., 2021).

### A.5 *Ex vivo* EAE data acquired at the Duke Center for In Vivo Microscopy (CIVM)

Magnetic resonance histology was performed on the cohort scanned at CIVM using a 9.4 T Agilent scanner solenoid coil using a radiofrequency (RF) refocused spin echo sequence. The reconstructed image size was 512 *×* 1024 *×* 512 with an isotropic resolution of 21.5 µm. For the purpose of the MacKenzie-Graham et al. (2009) study, the data were downsampled by a factor of 2 to produce a final image size of 256 *×* 512 *×* 256, yielding an isotropic resolution of 43 µm.

### A.6 *Ex vivo* EAE data acquired at the Beckman Institute at the California Institute of Technology

T2-weighted magnetic resonance microscopy images were acquired using an 11.7 T Bruker (Bruker Instruments, Billerica, MA) scanner. A RARE 3D imaging sequence was applied (matrix dimensions = 256 *×* 256 *×* 256; FOV = 3 cm *×* 1.5 cm *×* 1.5 cm; TR = 1500 ms; *TE*_eff_ = 10 ms; number of averages = 4) and the resulting image was zero-filled (Farrar & Becker, 2012; Fukushima, 2018) to yield a reconstructed image of approximately 60 µ*m*^3^. These images were then resampled to 50 µm *×* 50 µm *×* 47.5 µm resolution. For a more detailed description on these two *ex vivo* datasets, please see (MacKenzie-Graham et al., 2009).

The T1-weighted images were acquired using the same scanner (11.7 T Bruker) using the Fast Low Angle Shot (FLASH) sequence (matrix dimensions = 512 *×* 100 *×* 150; FOV = 3.3 cm *×* 1.1 cm *×* 1.65 cm; TR = 58.885 ms; TE = 7.713 ms; number of averages = 80).

### A.7 *Ex vivo* EAE data acquired at the Small Animal Imaging Core (SAIC) at Children’s Hospital Los Angeles

All *ex vivo* images collected at SAIC were acquired using an 11.7 T BioSpec (Bruker Instruments, Billerica, MA) scanner. The T1-weighted images were acquired using the following parameters: FLASH sequence; matrix dimensions = 600 *×* 300 *×* 300; FOV = 3 cm *×* 1.5 cm *×* 1.5 cm; TR = 23.3 ms; TE = 4.38 ms; number of averages = 24.

## B Processing and application of existing methods

The following summarizes the steps we performed to apply DeepBrainIPP and antsBrainExtraction on the test datasets for method comparison.

### B.1 DeepBrainIPP

During the application of DeepBrainIPP on our test datasets, we observed algorithm failures on a subset of our data. We surmised that these were because DeepBrainIPP could not accommodate some of our larger images. We addressed this by cropping the images as necessary to meet the DeepBrainIPP’s input size constraints before processing them with DeepBrainIPP. We cropped all *in vivo* isotropic test images with size 192 *×* 256 *×* 100 along the y-dimension to produce a cropped image of size 192 *×* 230 *×* 100. We similarly cropped four *ex vivo* test images, which were of size 256 *×* 512 *×* 256, to 256 *×* 460 *×* 256. All image cropping was done symmetrically, with an equal number of slices removed from both ends of the y-axis. We note that only slices not containing the brain were cropped.

We performed additional post-processing steps external to DeepBrainIPP’s workflow to consolidate its output brain labels. DeepBrainIPP performs two separate segmentations. First, it segments the larger brain area, which includes cerebrum, brainstem, and the cerebellum. DeepBrainIPP also performs a second segmentation to label to capture the smaller floccular nodules in the cerebellum called the bilateral paraflocculi, which is often missed in the larger brain mask. This method results in two separate segmentation files. In the DeepBrainIPP pipeline, the paraflocculus masks for *in vivo* and *ex vivo* data are resampled to 60 µm isotropic resolution, then zero-padded to produce an output mask of size 448 *×* 48 *×* 448. This is subsequently cropped to produce a final segmentation image of size 256 *×* 288 *×* 224. However, when segmenting the larger brain in the *in vivo* dataset, DeepBrainIPP resamples the data into 60 µm *×* 48 µm *×* 60 µm, and then zero pads the images to produce an output image of size 448 *×* 48 *×* 448. For *ex vivo* data, DeepBrainIPP resamples the image to 60 µm isotropic resolution and then crops it to produce an image size of 256 *×* 288 *×* 224. When cropping or padding, DeepBrainIPP first locates the image center by calculating the center of mass (COM) using image intensity values. The pipeline then adds or removes slices at the ends of the axes while preserving the COM (Alam et al., 2022).

As a result, the segmentations had different sizes and resolutions from each other as well as from the original input image. Moreover, some of the images were translated due to the cropping and padding. Therefore, we reversed all resampling and resizing processing steps to restore the masks to their original size and resolution. We used nearest-neighbor interpolation for resampling because DeepBrainIPP employs this method for resampling their training data and predicted labels. Lastly, we combined the brain and paraflocculus masks into a single mask by taking the union of the two masks.

### B.2 antsBrainExtraction

AntsBrainExtraction is an atlas-based method and requires a brain MRI template with the skull still attached and a corresponding probability map of the brain region. We generated six separate MRI templates and corresponding brain probability maps using the images from the training datasets. Two templates were rendered using *Chd8* and TBI training dataset. We created one template to represent the *in vivo* EAE, aging, and NeAt training datasets. We generated three additional templates using the *ex vivo* EAE dataset, which we subdivided into groups with similar MR contrasts. We created the templates using ANTs’ antsMultivariateTemplateConstruction2.sh script ^1^ (Avants et al., 2011). Using this script, we performed pairwise linear registration within each cohort to render four template images. Next, images were non-linearly registered to this template image and averaged to produce the final template image. The probability maps of the brain region were generated by applying the resulting transforms to the corresponding brain masks. Finally, antsBrainExtraction was applied to all of the images in the test dataset using the derived templates and brain priors. We note that in order to run antsBrainExtraction without error, we multiplied the pixel dimension sizes by 10 to approximate human MRI resolution.

## Footnotes

1 https://github.com/ANTsX/ANTs/blob/master/Scripts/antsMultivariateTemplateConstruction2.sh

